# Autophagy controls differentiation of *Drosophila* blood cells by regulating Notch levels in response to nutrient availability

**DOI:** 10.1101/2024.06.25.600418

**Authors:** Maximiliano J. Katz, Felipe Rodríguez, Fermín Evangelisti, Sebastián Perez-Pandolfo, Natalia Sommario, Agustina Borrat, Mariana Melani, Pablo Wappner

## Abstract

*Drosophila* larval hematopoiesis takes place at the lymph gland, where blood cell progenitors differentiate into two possible cell types: plasmatocytes, analogous to mammalian macrophages, or crystal cells that share features with mammalian megakaryocytes; a third cell type, the lamellocytes, can develop only upon specific immune challenges. In this work, we investigate the role of autophagy in *Drosophila* hematopoiesis. We found that autophagy inhibition in blood cell progenitors results in augmented crystal cell differentiation due to accumulation of high levels of Notch protein. Notch activation during hematopoiesis depends on the endocytic pathway, which cross-talks with autophagy: While endocytosis and endosomal maturation are essential for Notch activation, autophagosomes are required for Notch lysosomal degradation. TOR signaling inhibits autophagosome biogenesis, which in turn prevents the formation of Notch-containing amphisomes, being the latter necessary for Notch lysosomal destruction. Reduction of Notch lysosomal degradation shifts the balance towards Notch activation at late endosomal membranes, thereby enhancing differentiation of crystal cells. Our work defines a novel mechanism of regulation of immune cell differentiation in response to the nutritional status of the organism: High nutrient availability induces TOR activation, thereby inhibiting autophagy, hindering lysosomal degradation of Notch, and promoting crystal cell differentiation.

## Introduction

*Drosophila* larval hematopoiesis takes place at the lymph gland, where centrally located blood cell progenitors undergo differentiation to give rise to myeloid-like lineage cells, which include plasmatocytes and crystal cells (CCs), while a third cell-type, the lamellocytes, differentiates only upon certain specific immune challenges, such as parasitic wasp egg invasion (Cho et al., 2020; Girard et al., 2021). Plasmatocytes are macrophage-like cells that represent around 95% of total circulating blood cells, while CCs account for the remaining 5% (Banerjee et al., 2019) CCs display large cytoplasmic prophenoloxidase crystalline inclusions, which breakdown after the immune insult, releasing phenoloxidases to the blood stream, thus contributing to melanization and destruction of the pathogen (Binggeli et al., 2014).

The balance between maintenance and differentiation of blood cell progenitors of the lymph gland is tightly controlled by a wide array of factors ranging from signaling proteins (Benmimoun et al., 2012; Morin-Poulard et al., 2016; Terriente-Félix et al., 2017; Goyal et al., 2021; Kapoor et al., 2022) and metabolites (Dragojlovic-Munther and Martinez-Agosto, 2012; Goyal et al., 2021), to chemical and mechanical cues (Cho et al., 2018; Tian et al., 2023), which ultimately determine the progenitor fate. Particularly, differentiation of progenitors into CCs depends largely on the Notch pathway, activated, at least in part, by its ligand Serrate (Duvic et al., 2002; Lebestky et al., 2003; Mukherjee et al., 2011; Cho et al., 2018). Notch pathway inhibition in differentiation-competent blood cell progenitors provokes reduction or complete loss of CCs, while genetic overactivation of the pathway leads to excess of CCs (Lebestky et al., 2000; Blanco-Obregon et al., 2020).

Macroautophagy (referred hereafter as autophagy) is an evolutionary conserved self-degradative process, essential for housekeeping functions, such as recycling of macromolecules and organelles, as well as for removing protein aggregates. During autophagy, the cytoplasmic components to be degraded are sequestered in a double membrane organelle called autophagosome, which then fuses with lysosomes, where hydrolytic enzymes execute degradation of the cargo (Cuervo, 2004; Yorimitsu and Klionsky, 2005; Feng et al., 2014; Nakatogawa, 2020). Alternatively, an intermediate step may take place, as autophagosomes can fuse with late endosomes/multivesicular bodies to generate amphisomes, which in turn fuse with lysosomes, where degradation of the cargo takes place (Berg et al., 1998; Fader et al., 2008; Nakamura and Yoshimori, 2017). Although autophagy is essentially a bulk degradation process, selectivity is attained by autophagy adaptor proteins that bind to certain cargoes, such as organelles and aggregation-prone misfolded proteins or protein aggregates (Bjørkøy et al., 2005; Anding and Baehrecke, 2017).

In spite of autophagy being an essentially housekeeping recycling process, it is of fundamental importance in various cell differentiation events across eukaryotic organisms (Vázquez et al., 2012; Zhang et al., 2012; Wang et al., 2014a; Chen et al., 2018; Bankston et al., 2019; Clarke and Simon, 2019; Varga et al., 2022; Tan et al., 2024). Particularly, in mammalian hematopoiesis, loss-of-function experiments have revealed that autophagy participates in differentiation or maintenance of most immune cell types from both the myeloid and lymphoid lineages (Jacquel et al., 2012; Riffelmacher et al., 2017; Riffelmacher et al., 2018; Menshikov et al., 2021; Metur and Klionsky, 2021). However, the mechanisms by which autophagy regulates hematopoiesis are poorly defined at the molecular and cellular level.

In the current study, we show that basal autophagy in the *Drosophila* lymph gland is particularly active, and that restrains differentiation of CCs. Inhibition of autophagy in blood cell progenitors results in increased levels of Notch, whose regulation depends largely on the endocytic pathway: While endocytosis and endosome maturation are necessary for Notch activation, formation of multivesicular body vesicles, and fusion of the multivesicular body with lysosomes are required for Notch lysosomal degradation. Notch lysosomal degradation is regulated by autophagy through amphisome formation, and autophagy is in turn regulated by nutrient availability through the TOR pathway. Our study therefore establishes a mechanistic link between the nutritional status of the organism and blood cell differentiation through the regulation of autophagy and Notch activation.

## Results

### High levels of basal autophagy in the lymph gland are required for normal hematopoiesis

To investigate the role that autophagy plays in blood cell differentiation in the lymph gland of third instar larvae, we first analyzed the levels of basal autophagy in this organ. Two different criteria, namely nucleation of the autophagosome reporter 3xmCherry-Atg8a and Lysotracker staining (Hegedűs et al., 2016), revealed that basal autophagy levels are particularly high at the lymph gland in well-fed conditions, as compared with other *Drosophila* larval tissues (SFig 1A-G). As in other biological settings, autophagy was further activated in the lymph gland when larvae were subjected to a 6h starvation (SFig 1H-J). Canonical autophagy pathway regulators were required for basal autophagy in the lymph gland, as indicated by knock-down of different components of the autophagy pathway (Chang and Neufeld, 2009) (SFig 2). These data suggest that the canonical autophagy pathway is active at unusually high levels in the lymph gland, as compared with other *Drosophila* organs.

To investigate a possible role of autophagy in the differentiation of blood cells, we performed loss-of-function experiments of autophagy pathway genes and analyzed blood cell specific markers. In *Atg1^Δ3D^* homozygous mutant larvae, autophagy was suppressed (Fig 1A) and crystal cells increased dramatically (Fig 1D, E), while the proportion of blood cell progenitors and plasmatocytes were not significantly modified (Fig 1B, C).

**Figure 1.**
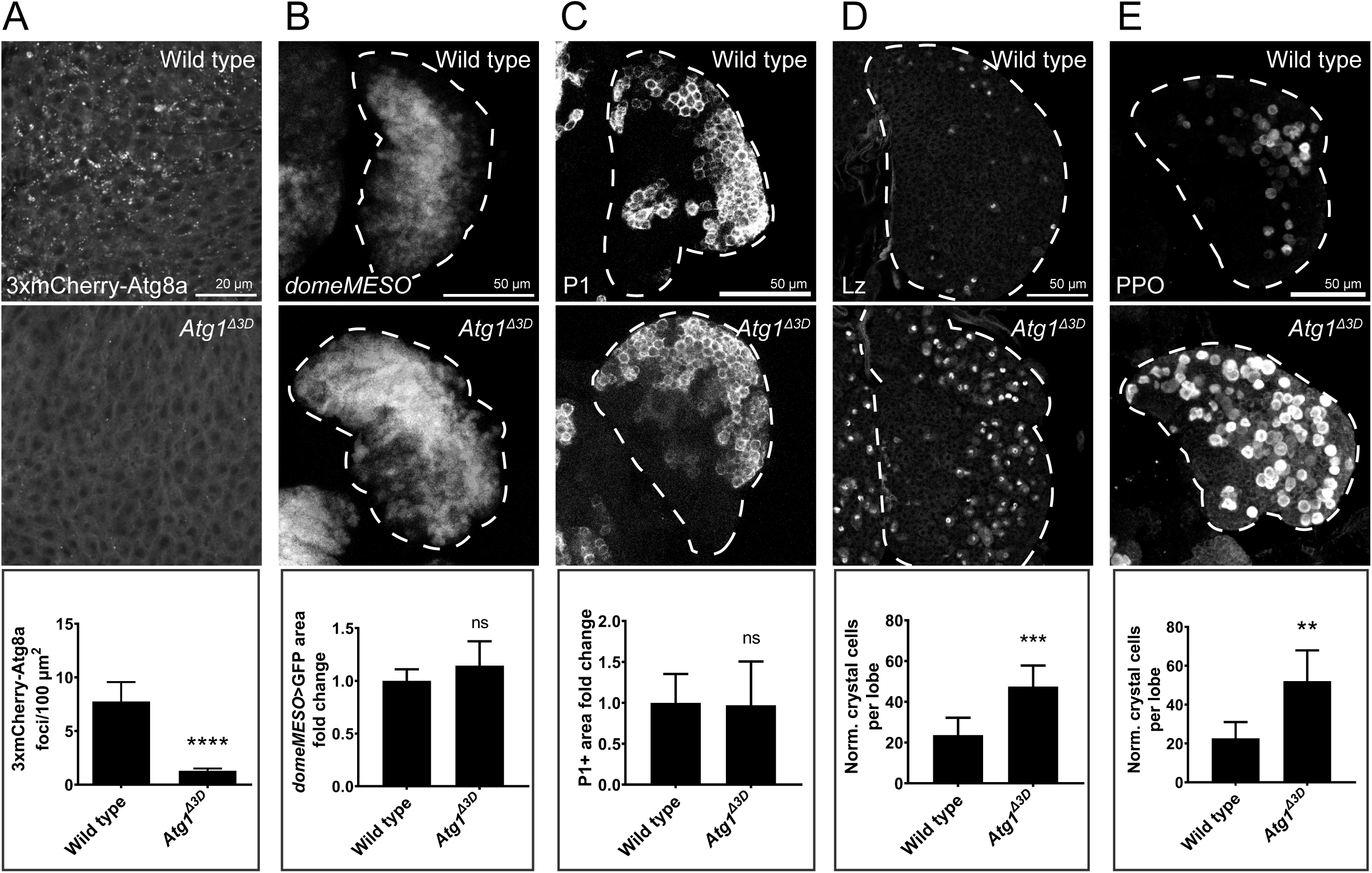
Crystal cell differentiation increased in *Atg1^Δ3D^* mutants. Confocal images of individual lobes of larval lymph glands (white dashed lines). In *Atg1^Δ3D^* homozygous mutants, autophagy activation was reduced, as indicated by 3xmCherry-Atg8a nucleation (A; n ≥ 8 primary lobes). Progenitor (*domeMESO*>GFP; B; n ≥ 13) or plasmatocyte (P1; C; n ≥ 24) populations were unaffected in *Atg1* mutants. Crystal cells increased in *Atg1^Δ3D^* mutant larvae as assessed by either anti-Lozenge (Lz; D; n ≥ 7) or anti-prophenoloxidase (PPO; E; n ≥ 10). Data are represented as mean ± 95% CI and were assessed by Student’s *t*-test. **p<0.01, ***p<0.001, ****p<0.0001, ns, not significant, p>0.05.

Given the increase of crystal cell differentiation in the *Atg1^Δ3D^* mutant lymph gland, we next wandered if autophagy is required cell-autonomously in blood cell progenitors for normal crystal cell differentiation, so we expressed double stranded RNAs against various autophagy regulators specifically in this cell type by using a *domeMESO*-Gal4 driver. Knock-down (KD) of Atg1 or Atg17 (components of the autophagy initiation complex) (Fig 2A, B, C), of Vps15 or Vps34 (components of the nucleation complex) (Fig 2A, D, E), or of Atg18 (involved in phagophore expansion) (Nakatogawa, 2020) (Fig 2A, F) provoked clear increase of CC differentiation (Fig 2G, H), as visualized with anti-Lozenge (Lz) or anti-prophenoloxidase (PPO) antibodies (Mukherjee et al., 2011; Blanco-Obregon et al., 2020). These results indicate that autophagy is required in blood cell progenitors to restrain CC differentiation.

**Figure 2.**
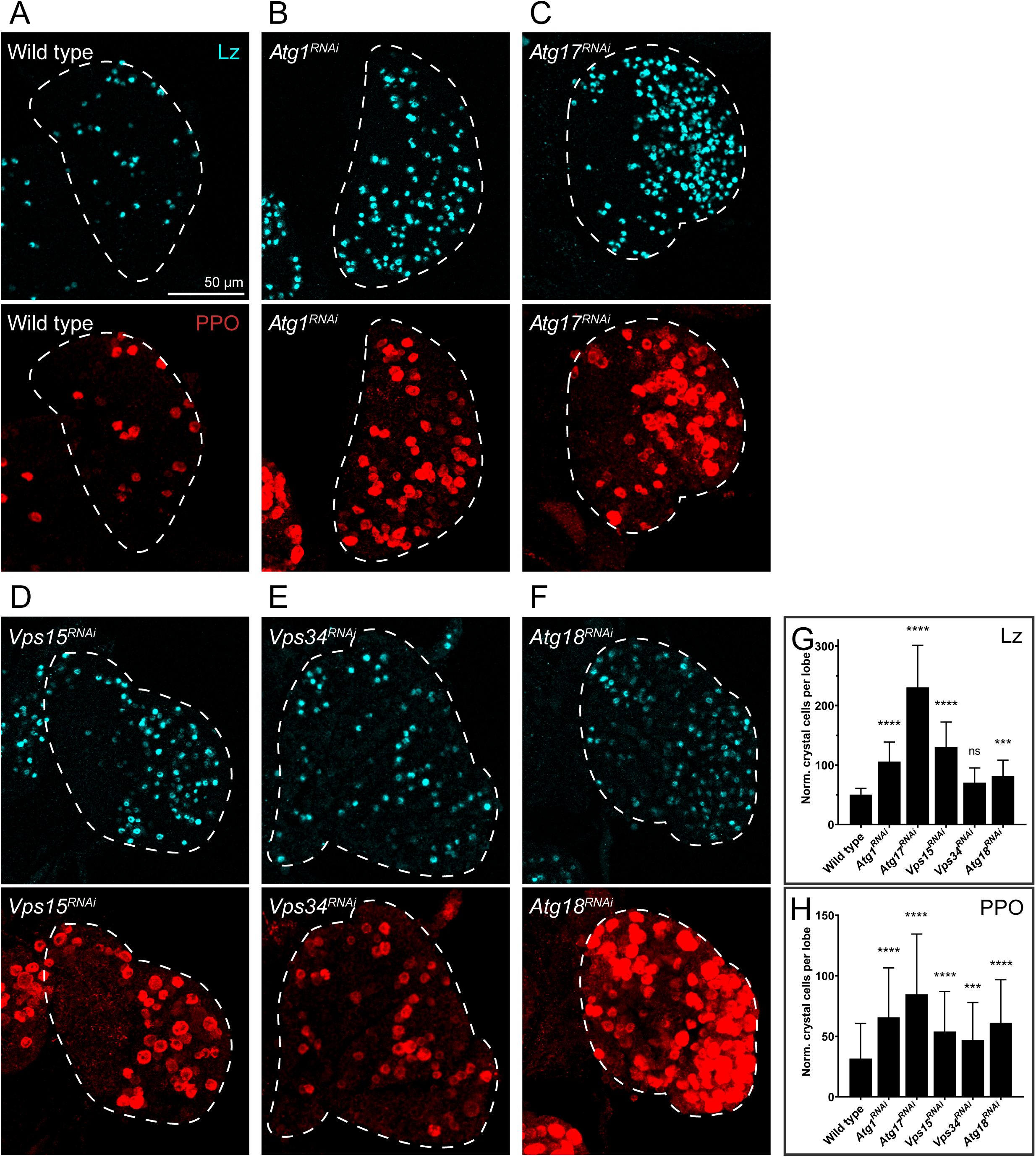
Knock-down of autophagy pathway genes in blood cell progenitors provoked increased crystal cell differentiation. Crystal cells were visualized by anti-Lozenge (Lz) or anti-prophenoloxidase (PPO) immunofluorescence in lymph gland lobes where the indicated double-stranded RNAs, affecting different autophagy genes, have been expressed under control of a *domeMESO*-Gal4 driver (A-F). Crystal cell differentiation increased in comparison to wild type controls (*UAS-LacZ*). Quantification of the normalized number of crystal cells per lobe is depicted on panels G (anti-Lz) and H (anti-PPO), where data are represented as mean ± 95% CI. Statistical analysis employed was a Likelihood Ratio Test followed by Dunnett’s test for multiple comparisons (treatments versus control). ***p<0.001, ****p<0.0001, ns, not significant, p>0.05. For wild type, n = 65 primary lobes; for *Atg1^RNAi^*, n = 24; for *Atg17^RNAi^*, n = 25; for *Vps15^RNAi^*, n = 19; for *Vps34^RNAi^*, n = 13; for *Atg18^RNAi^*, n = 21 for Lz, n = 20 for PPO.

### Autophagy sets a limit to Notch pathway activation and crystal cell differentiation

It has been shown that differentiation of CCs largely depends on activation of the Notch pathway, stimulated at least in part by its ligand Serrate (Lebestky et al., 2000; Duvic et al., 2002; Lebestky et al., 2003; Mukherjee et al., 2011; Blanco-Obregon et al., 2020). We analyzed whether ligand-independent activation of Notch (Hounjet and Vooijs, 2021) also contributes to CC differentiation in the lymph gland. This seems to be the case, as progenitor-specific overexpression of the E3 ubiquitin ligase deltex (dx), that promotes ligand-independent Notch endocytosis and Notch pathway activation (Matsuno et al., 1995; Yamada et al., 2011; Shimizu et al., 2024), provoked significant increase of CC differentiation (SFig 3A, B, D). Consistent with this, CC differentiation was also enhanced by the KD of Suppressor of deltex (Su(dx)), another E3 ligase that in this case inhibits non-canonical Notch pathway activation by promoting Notch lysosomal degradation (Fostier et al., 1998; Cornell et al., 1999) (SFig 3A, C, D). These observations support the notion that ligand-independent Notch pathway activation contributes to CC differentiation in the lymph gland.

Next, we conducted genetic interaction experiments to explore if autophagy-dependent regulation of Notch may account for the effect of autophagy on CC differentiation. As expected, inhibition of autophagy in progenitors failed to promote CC differentiation when Notch expression was simultaneously silenced (Fig 3A-D, I). Interestingly, the increase of CCs observed upon KD of Atg1 was suppressed in Notch (Fig 3A, B, E, F, I) or Suppressor of Hairless (Fig 3A, B, G-I) heterozygous mutant larvae, suggesting that autophagy antagonizes Notch activation during CC differentiation. Similarly, genes that participate of the non-canonical (ligand-independent) Notch pathway also showed antagonistic genetic interactions with autophagy: KD of dx, which on itself does not modify CC differentiation, suppressed the enhancement of CC differentiation induced by KD of Atg1 (SFig 4A-D, G). Likewise, overexpression of Su(dx), which on itself has no effect on differentiation, suppressed the enhancement of CCs provoked by Atg1 silencing (SFig 4A, B, E-G). Taken together, these results suggest that autophagy sets a limit to Notch pathway activation in the lymph gland, thereby restraining CC differentiation.

**Figure 3.**
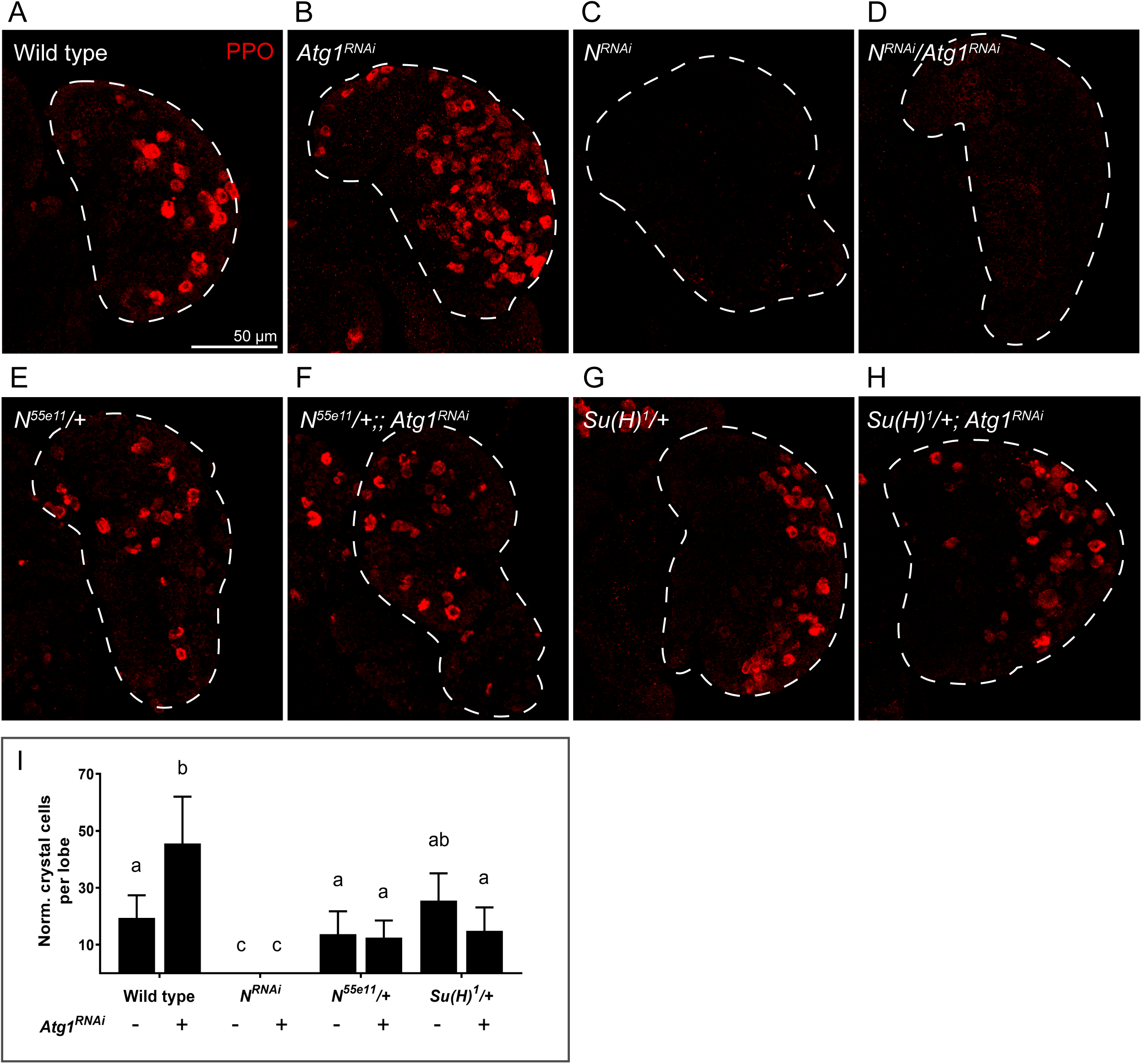
Antagonistic genetic interactions between autophagy and Notch pathway genes. Crystal cells were visualized in Z-stack projections of lymph glands single lobes by anti-PPO immunofluorescence. Expression of a *Notch* RNAi, driven by *domeMESO*-Gal4, completely blocked crystal cell differentiation (A, C, D), while expression of an *Atg1* RNAi increased crystal cell number (A, B). The increase of crystal cells provoked by *Atg1* RNAi expression was suppressed in *Notch* (*N^55e11^*, E, F) or *Suppressor of Hairless* (*Su(H)^1^*, G, H) heterozygous mutant larvae. In *N^55e11^* (E) or *Su(H)^1^* (G) heterozygous larvae not expressing any RNAi, the number of crystal cells was normal. (I) Quantification of the normalized number of crystal cells per lymph gland lobe. Data represented as mean ± 95% CI. Statistical analysis performed was Kruskal-Wallis test followed by Dunn’s test for multiple comparisons. Distinct letters represent statistically significant differences (p<0.05). For wild type, n = 33 primary lobes; for *Atg1^RNAi^*, n = 43; for *N^RNAi^*, n = 18; for *N^RNAi^/Atg1^RNAi^*, n = 23; for *N^55e11^/+*, n = 14; for *N^55e11^/+; Atg1^RNAi^*, n = 28, for *Su(H)^1^/+*, n = 18; for *Su(H)^1^/+; Atg1^RNAi^*, n = 18.

Then, we investigated if the effect of autophagy on Notch pathway activation may rely on direct control of Notch receptor abundance. This was indeed the case, as KD of autophagy genes in blood cell progenitors provoked significant increase of Notch protein levels, as revealed by anti-Notch immunofluorescence (Fig 4A-F). Downregulation of autophagy also provoked increased Notch-dependent transcription, indicated by the activity of an Enhancer of Split transcriptional reporter (Rodrigues et al., 2021) (Fig 4A-E, G). These results indicate that upon autophagy inhibition, Notch protein accumulates, and the Notch pathway is overactivated.

**Figure 4.**
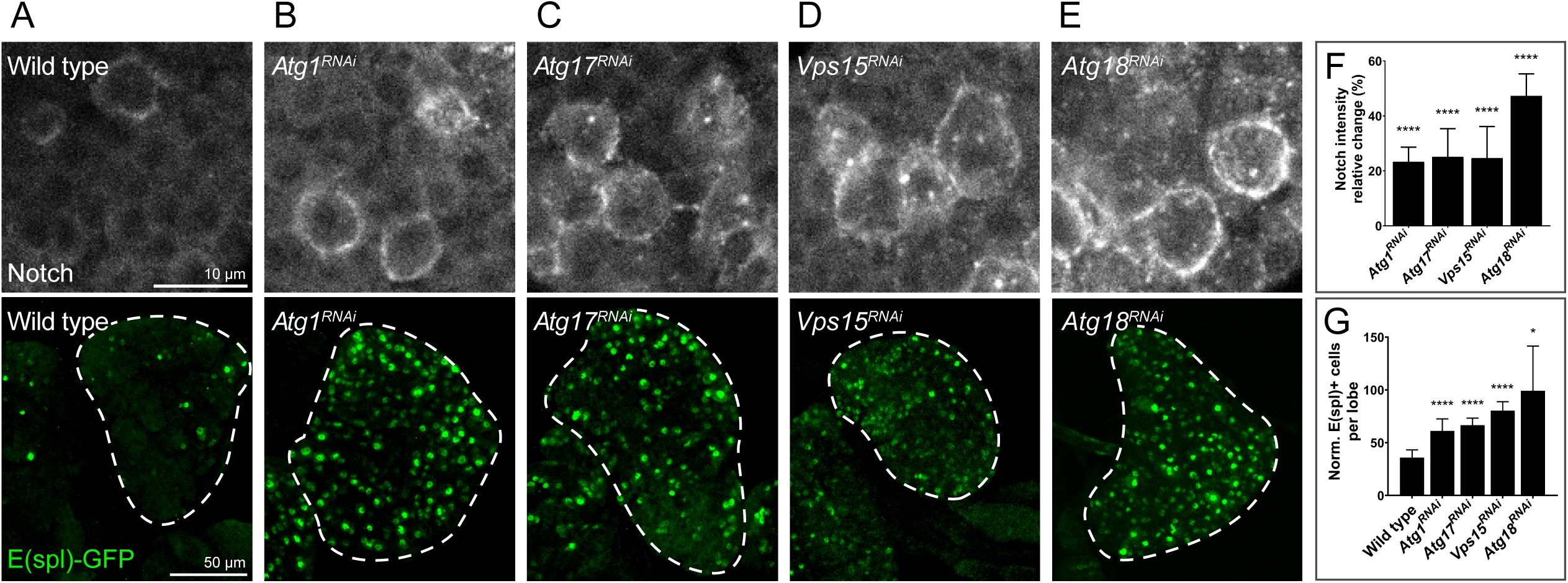
Notch protein levels and Notch-dependent transcription increased upon knock-down of autophagy genes in blood cell progenitors. Anti-Notch immunofluorescence revealed that Notch levels increased significantly in lymph gland lobes where double-stranded RNAs against the indicated autophagy genes (B-E) were expressed with a *domeMESO*-Gal4 driver, in comparison to wild type control individuals (A, *UAS-LacZ*). Mean fluorescence intensity was quantified in single slices, and relative change of immunofluorescence respect to the control was calculated and plotted as mean ± 95% CI (F). For wild type, n = 37 primary lobes; for *Atg1^RNAi^*, n = 25; for *Atg17^RNAi^*, n = 16; for *Vps15^RNAi^*, n = 16; for *Atg18^RNAi^*, n = 25. Notch transcriptional activity was evaluated with an E(spl)-GFP transcriptional reporter in each genotype (A-E), showing that the increase of Notch protein levels was paralleled by Notch-dependent transcription. (G) Quantification of the normalized number of GFP-positive cells per lobe represented as mean ± 95% CI. For wild type, n = 42 primary lobes; for *Atg1^RNAi^*, n = 21; for *Atg17^RNAi^*, n = 20; for *Vps15^RNAi^*, n = 55; for *Atg18^RNAi^*, n = 13. Statistical analysis (F, G) were performed using one-way ANOVA followed by Dunnett’s test for comparisons of treatments versus control. *p<0.05, ****p<0.0001.

### Cross-talk between the endocytic and autophagy pathways accounts for autophagy-dependent regulation of Notch and crystal cell differentiation

We next wandered about the mechanism by which autophagy controls Notch abundance. Autophagy is essentially a bulk degradation process and, with few exceptions (Gao et al., 2010; Mancias et al., 2014; Simpson et al., 2024), does not usually target specific signaling molecules. We therefore reasoned that a crosstalk between the autophagy machinery and the endocytic pathway -essential for Notch signaling in some biological contexts (Le Borgne et al., 2005; Hounjet and Vooijs, 2021)- might account for autophagy-dependent Notch degradation. In the canonical pathway, after binding its ligand, the Notch receptor is cleaved at the plasma membrane first by the metalloprotease Kuzbanian (Kuz), and then by the γ-secretase complex, releasing the Notch Intracellular Domain (NICD) that enters the nucleus and controls gene expression (Fig 5A) (Bray, 2016). However, in some biological contexts, Notch is activated by the γ-secretase not only at the plasma membrane but also at the membrane of endosomes (Fig 5B): after being cleaved by Kuz, Notch undergoes endocytosis, ending up in the membrane of early endosomes (EEs), which then acidify and mature to late endosomes (LEs), and finally evolve into multivesicular bodies (MVBs) (Huotari and Helenius, 2011). In this context, the γ-secretase can cleave Notch either at the membrane of LEs, or at the limiting membrane of MVBs, releasing the NICD that enters the nucleus and regulates transcription (Fig 5B) (Vaccari et al., 2008). This mechanism of Notch activation can also take place in a ligand-independent manner (Fig 5C), in which case full-length Notch undergoes endocytosis without being cleaved by Kuz, and then the same process as in the canonical (ligand-dependent) Notch pathway takes place (Fig 5C) (Schnute et al., 2018).

**Figure 5.**
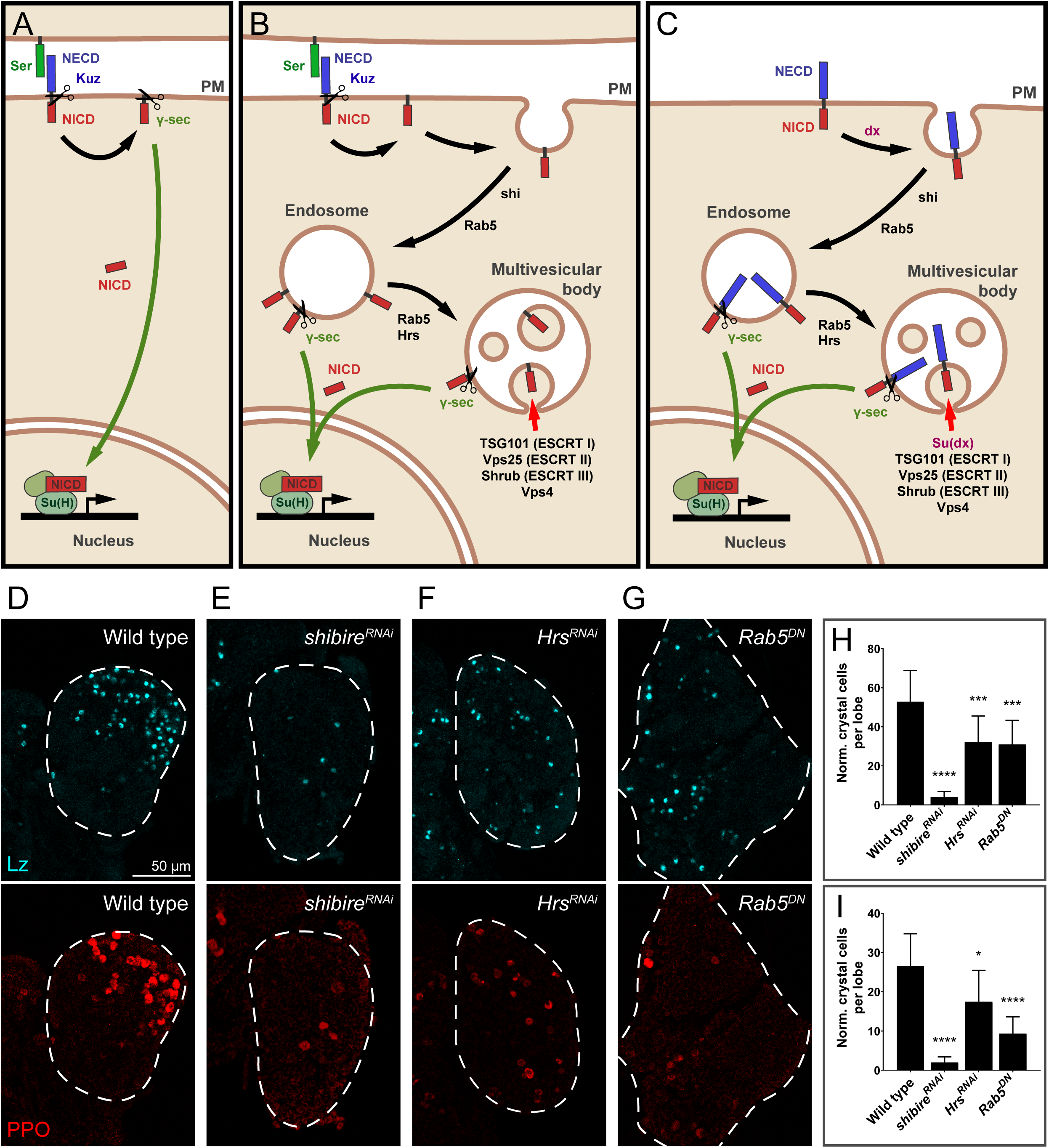
The endocytic pathway controls crystal cell differentiation. (A, B, C) Schematic representation of the Notch pathway. (A) Upon binding of a ligand (Serrate (Ser) in this case), Notch undergoes cleavage at the plasma membrane mediated first by the metalloprotease Kuzbanian (Kuz) and then by the γ-secretase complex (γ-sec). After cleavage, the Notch Intracellular Domain (NICD) is released, enters the nucleus and regulates gene expression together with other transcriptional coactivators that include Suppressor of Hairless (Su(H)). (B) Alternatively, the second cleavage of Notch, mediated by γ-sec, can take place in endosomal membranes: after Ser binding, and cleavage by Kuz, Notch undergoes endocytosis, which is dependent on shibire (shi) among others, and remains inserted in the membrane of early endosomes that will mature by the action of Hrs and Rab5 to become late endosomes. Invagination of the limiting membrane of late endosomes, mediated by genes of the ESCRT complexes 0-III will give rise to the multivesicular body. Some of the Notch molecules remain at the limiting membrane of the multivesicular body, while others end up in the membrane of intraluminal vesicles. The proteins Tsg101, Vps25 and Shrub are part of the ESCRT complexes I, II and III respectively. Vps4 is the effector ATPase of the ESCRT complexes. The Notch domain present at the limiting membrane of late endosomes or multivesicular bodies can be cleaved by the γ-sec to release the NICD that enters nucleus to regulate transcription. The Notch domains present in intraluminal vesicles of the multivesicular body are degraded at the lysosome (not shown in the scheme). (C) Ligand-independent activation of Notch: Notch can undergo endocytosis even in the absence of ligand binding; in this case, the entire (uncleaved) receptor ends up in the membrane of endosomes. The subsequent steps of the pathway are identical to those described in (B). Ligand-independent endocytosis of Notch is stimulated by the E3 ubiquitin ligase deltex (dx); formation of Notch-containing intraluminal vesicles of the multivesicular body, and subsequent lysosomal degradation (not shown in the scheme), is stimulated by the activity of the E3 ubiquitin ligase Suppressor of deltex (Su(dx)). Crystal cell differentiation, assessed by anti-Lozenge (Lz) or anti-prophenoloxidase (PPO) immunofluorescence, was reduced after knock-down of the dynamin shibire (E), or of Hrs (F) or Rab5 (G), with a *domeMESO*-Gal4 driver. (H, I) Quantification of the results shown in panels (D-G) represented as mean value of normalized crystal cells per lobe ± 95% CI. For wild type, n = 86 primary lobes; for *shibire^RNAi^*, n = 24; for *Hrs^RNAi^*, n = 24; for *Rab5^DN^*, n = 22. The statistical analysis performed was a Likelihood Ratio Test followed by Dunnett’s test for comparisons of treatments against control. *p<0.05, ***p<0.001, ****p<0.0001.

We therefore explored whether normal progression of the endocytic pathway is required for Notch activation and CC differentiation in the lymph gland, and then, if the endocytic pathway may crosstalk with autophagy, thereby accounting for autophagy-dependent regulation of Notch (Fig 4). Indeed, blood cell progenitor-specific KD of shibire, a dynamin essential for completing endocytosis (van der Bliek and Meyerowrtz, 1991), decreased CC differentiation (Fig 5D, E, H, I), suggesting that endocytosis is required for Notch activation in this context. Likewise, KD of hrs, required for EE to evolve into LE, or expression of a dominant-negative form of the small GTPase Rab5, required for EE formation and maturation (Bucci et al., 1992; Vaccari et al., 2008), both significantly reduced CC differentiation (Fig 5D, F-I).

Invagination of the limiting membrane of the MVB, leading to intraluminal vesicle formation (Fig 5B,C), is promoted by four ESCRT complexes (Endosomal Sorting Complex Required for Transport 0 to III) (Katzmann et al., 2001; Hurley, 2015), and incorporation of Notch to intra luminal vesicles results in Notch lysosomal degradation (Jia et al., 2009; Yamamoto et al., 2010). To gather further evidence that the endocytic pathway is necessary for Notch regulation, and particularly to assess if ESCRT complexes may promote Notch degradation in blood cell progenitors, we silenced the expression of Tsg101, a component of ESCRT-I complex, of Vps25, component of ESCRT-II, of Shrub, member of ESCRT complex III, or the expression of Vps4, the effector ATPase of all ESCRT complexes (Williams and Urbé, 2007; Vietri et al., 2020) (Fig 5B, C). Silencing of any of the above genes, expected to inhibit intraluminal vesicle formation and Notch lysosomal degradation, provoked clear enhancement of CC differentiation (Fig 6A-G), and increased Notch protein levels (Fig 6A-E, H), further supporting the notion that the endocytic pathway regulates Notch in blood cell progenitors, thereby controlling CC differentiation.

**Figure 6.**
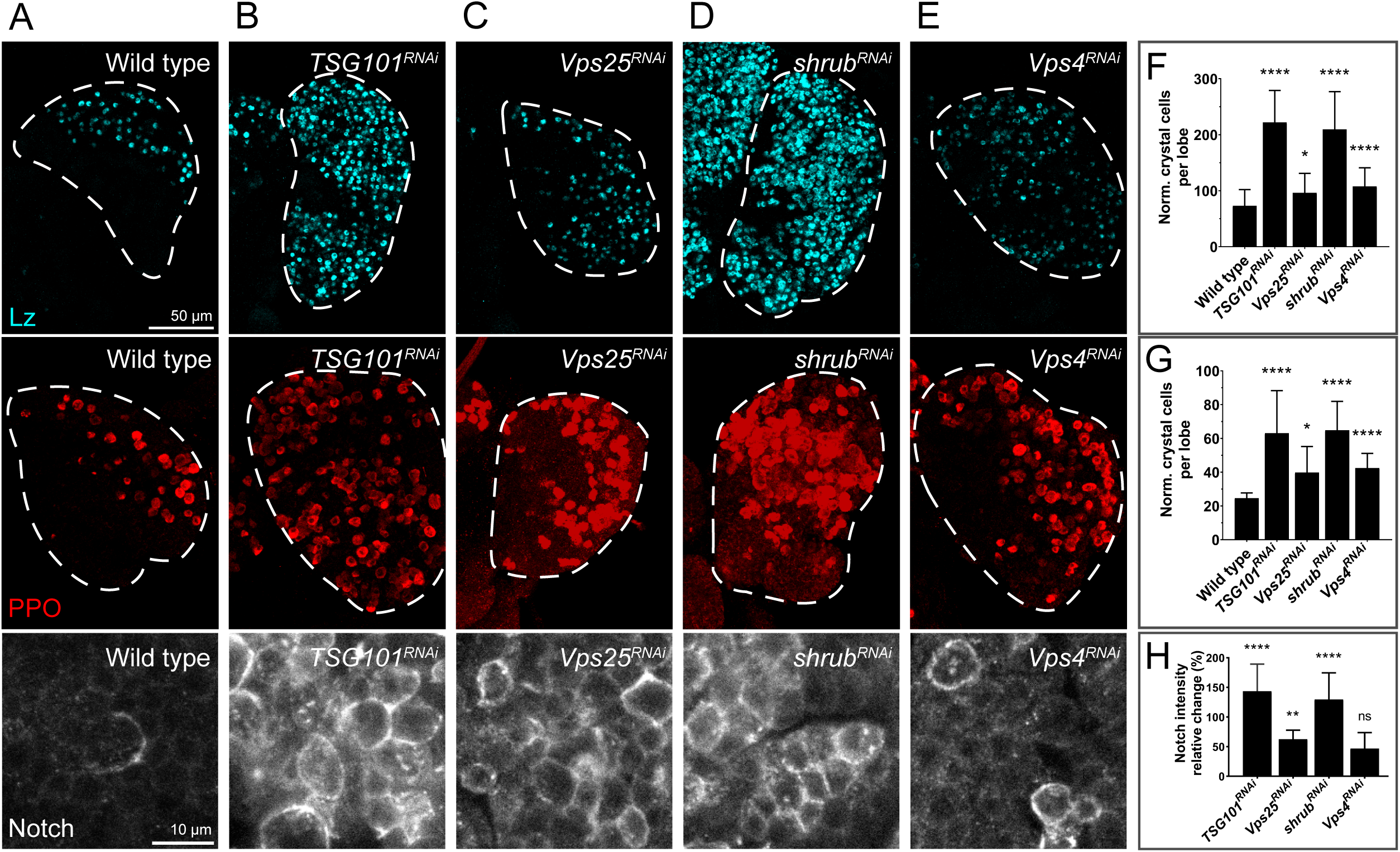
Knock-down of elements of ESCRT complexes resulted in increased Notch abundance and enhanced crystal cell differentiation. Anti-Lozenge (Lz) and anti-prophenoloxidase (PPO) stainings to visualize crystal cells, and assess the effect of *domeMESO*-Gal4-driven knock-down of genes of different ESCRT complexes (B-D) or the gene *Vps4* (E) that encodes the effector ATPase of ESCRT complexes, in comparison with wild type individuals (A). (F, G) Quantification of normalized crystal cells per lobe as assessed by anti-Lz or anti-PPO staining respectively, visualized as mean ± 95% CI. One-way ANOVA followed by Dunnett’s test for multiple comparisons (treatments versus control). For wild type, n = 86 primary lobes; for *TSG101^RNAi^*, n = 24 for Lz and n = 22 for PPO; for *Vps25^RNAi^*, n = 24; for *shrub^RNAi^*, n = 19; for *Vps4^RNAi^*, n = 33. Notch immunofluorescence performed with an antibody targeting the Notch extracellular domain shows that Notch protein levels in each genotype (A-E) parallels the increase in differentiation of crystal cells. (H) Quantification of Notch immunofluorescence expressed as relative change in fluorescence intensity respect of the wild type genotype plotted as mean ± 95% CI. For wild type, n = 20; for *TSG101^RNAi^*, n =16; for *Vps25^RNAi^*, n = 19; for *shrub^RNAi^*, n = 15; for *Vps4^RNAi^*, n = 14. One-way ANOVA followed by Dunnett’s test for multiple comparisons (treatments versus control). *p<0.05, **p<0.01, ****p<0.0001, ns, not significant (p>0.05).

Having established that the endocytic pathway plays a central role in Notch regulation in blood cell progenitors during CC differentiation, we next wondered whether autophagy impinge on Notch activation through the endocytic pathway. It is well known that autophagosomes can fuse with MVBs to generate amphisomes, and that amphisomes in turn fuse with lysosomes to generate autolysosomes (Berg et al., 1998; Zhao et al., 2021) (SFig 5A). Moreover, it has been proposed that autolysosomes, where degradation of the autophagosome cargo takes place (Zhao et al., 2021), are formed exclusively through this intermediate fusion event (Filimonenko et al., 2007; Razi et al., 2009). We therefore investigated if inhibition of autophagosome formation may lead to accumulation of MVBs, stalling Notch lysosomal degradation, and shifting the balance towards Notch activation. In agreement with this hypothesis, super-resolution confocal imaging (AiryScan) revealed that in wild type larvae, Notch appeared within or associated with Rab5-positive vesicles (SFig 5B), or Rab7-positive vesicles (SFig 5C, D), which correspond respectively to EE, and LE/MVBs (Behnia and Munro, 2005; Dunst et al., 2015). Upon autophagy inhibition in blood cell progenitors, Rab7-positive vesicles (LE/MVBs) significantly enlarged (SFig 5C-E), and contained increased amounts of Notch protein (SFig 5C, D, F), supporting the notion that autophagy mediates lysosomal degradation of Notch that is present in endosomes. Furthermore, in wild type individuals Notch colocalized with the autophagosomal marker Atg8a and/or with the endolysosomal marker Lamp (Akbar et al., 2009) or with Rab7 (SFig 6), suggesting that Notch traffics through amphisomes and autolysosomes prior to lysosomal degradation. Taken together, our results support a model in which autophagy regulation, through the fusion of autophagosomes with MVBs, sets a balance between Notch activation at the limiting membrane of LE/MVBs, and Notch lysosomal degradation once it has been incorporated to intraluminal vesicles of the MVB. Consistent with such model, silencing of the phosphatidylinositol(3)P-5-kinase Fab1, required for endolysosomal maturation and function (Rusten et al., 2006) (SFig 5A), enhanced CC differentiation (Fig 7A, B, E, F). Likewise, silencing of the SNAREs Vamp7 or Syntaxin-17, required for fusion of autophagosomes with MVBs as well as fusion of autophagosomes or amphisomes with lysosomes (Itakura et al., 2012; Takáts et al., 2013) (SFig 5A), enhanced CC differentiation (Fig 7A, C-F). These results suggest that prevention of autolysosome formation inhibits Notch lysosomal degradation, thus favoring instead Notch activation at the limiting membrane of LEs/MVBs.

**Figure 7.**
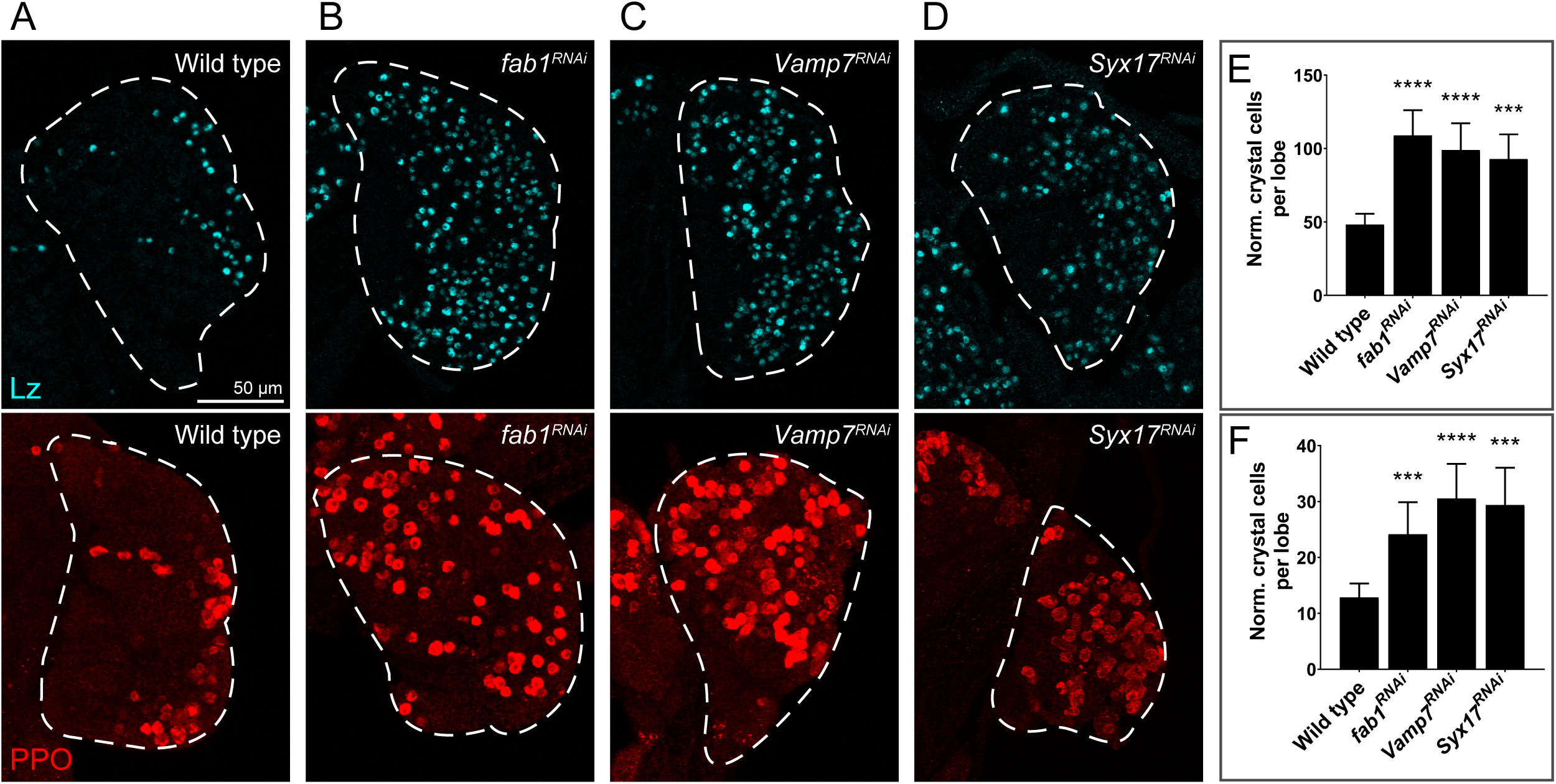
Blockage of lysosomal maturation or impairment of fusion events that give rise to amphisomes or autolysosomes increased crystal cell differentiation. Anti-Lozenge (Lz) and anti-prophenoloxidase (PPO) stainings to visualize crystal cells. After knock-down of Fab1 with a *domeMESO*-Gal4 driver, an increase of crystal cell differentiation occurred in comparison with wild type control larvae (A, B). Knock-down of the SNAREs Vamp7 or Syntaxin-17 also led to augmented crystal cell differentiation (C, D). (E, F) Quantification of normalized crystal cells per lobe, as assessed with anti-Lz and anti-PPO antibodies respectively. Mean ± 95% CI. Wild type, n = 22; *fab1^RNAi^*, n = 21; *Vamp7^RNAi^*, n = 20; *Syx17^RNAi^*, n = 19. Brown-Forsythe and Welch ANOVA followed by Dunnett’s T3 test for comparisons of treatments against control. ***p<0.001, ****p<0.0001.

### Nutrient availability regulates autophagy through the TOR pathway, thereby controlling crystal cell differentiation

In eukaryotic cells, nutrient availability mediates regulation of autophagy through the TOR pathway (TOR) (Scott et al., 2004; Neufeld, 2010; Kim et al., 2011), so we sought to assess if alterations in the composition of the larval culture medium impinge on CC differentiation by modulating the autophagy-Notch axis in blood cell progenitors. After increasing the amount of yeast in the medium, which augments amino acid availability (Lee et al., 2008a), autophagy was significantly reduced (SFig 7A, B), while Notch abundance increased significantly (Fig 8A, B), and CC differentiation was significantly enhanced (Fig 8C, G). The increase of CC differentiation in the yeast-rich medium was suppressed upon Atg1 overexpression (Fig 8D, G), a treatment that increases autophagy (Scott et al., 2007). Additionally, the increase of CCs in the yeast-rich medium (Fig 8C) was mitigated after knocking-down the amino acid transporter Slimfast (Slif) (Colombani et al., 2003) (Fig 8E, G). Conversely, upon overexpression of Slif in normal larval medium, strong increase of CC differentiation was observed (Fig 8C, F, G). Overall, these results indicate that an amino acid-rich diet inhibits autophagy in the lymph gland, leading to Notch accumulation, and increased crystal cell differentiation.

**Figure 8.**
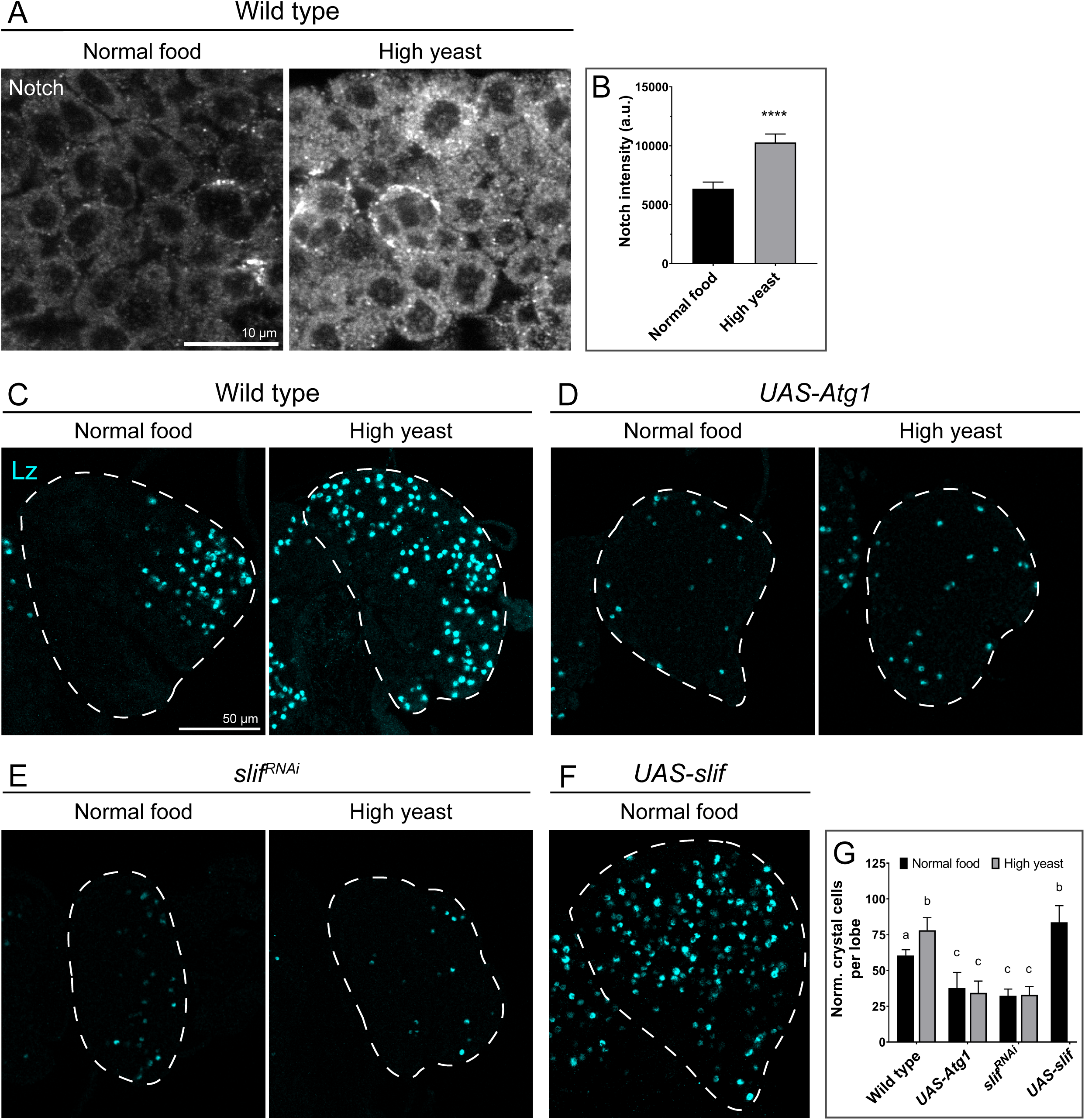
Amino acid availability controlled Notch protein abundance and crystal cell differentiation. Anti-Notch immunofluorescence (A) indicated that Notch protein levels were higher in lymph glands of larvae grown in culture medium containing high yeast (4% w/v) in comparison to larvae grown in normal medium (2% yeast w/v). (B) Mean Notch immunofluorescence intensity (a.u.) ± 95% CI, Student’s *t-*test (****, p<0.0001). For normal food, n = 13 primary lobes, for high yeast, n = 14. Crystal cell differentiation increased in larvae grown in the yeast-rich medium, in comparison to normal fly food (C), as assessed by anti-Lozenge (Lz) immunofluorescence. The increase of crystal cell differentiation observed in (C) was suppressed after overexpression of Atg1 (D) with *domeMESO*-Gal4 driver. RNAi-mediated knock-down of the amino acid transporter Slimfast (Slif) prevented the increase of crystal cell differentiation provoked by the addition of extra-yeast in the fly food (E). Overexpression of Slif led to increased crystal cell differentiation in larvae reared in normal food, compared with wild type animals (C, F). (G) Quantification of the results of panels (C-F), mean normalized crystal cells per lobe ± 95% CI. One-way ANOVA followed by Tukey’s test for multiple comparisons. Different letters represent statistically significant differences (p<0.05). For the wild type genotype in normal food, n = 144 primary lobes, and n = 51 for high yeast; for *UAS-Atg1*, n = 24 in normal food and n = 24 in high yeast; for *slif^RNAi^*, n = 75 in normal food and n = 44 in high yeast; for *UAS-slif,* n = 77.

The increase of CC differentiation observed in response to augmented nutrient availability was conveyed by the TOR pathway, since in *Rheb ^PΔ1^* or *TOR^2L19^* heterozygous loss-of-function mutants, CC differentiation was mitigated (Fig 9A-E). Consistent with this, overexpression of Akt or Rheb, as well as KD of Tsc1, treatments that inhibited autophagy (SFig 7D-H), significantly enhanced CC differentiation (Fig 9F-K). These results suggest that amino acid availability regulates activation of the TOR pathway in blood cell progenitors, thereby controlling autophagy and crystal cell differentiation.

**Figure 9.**
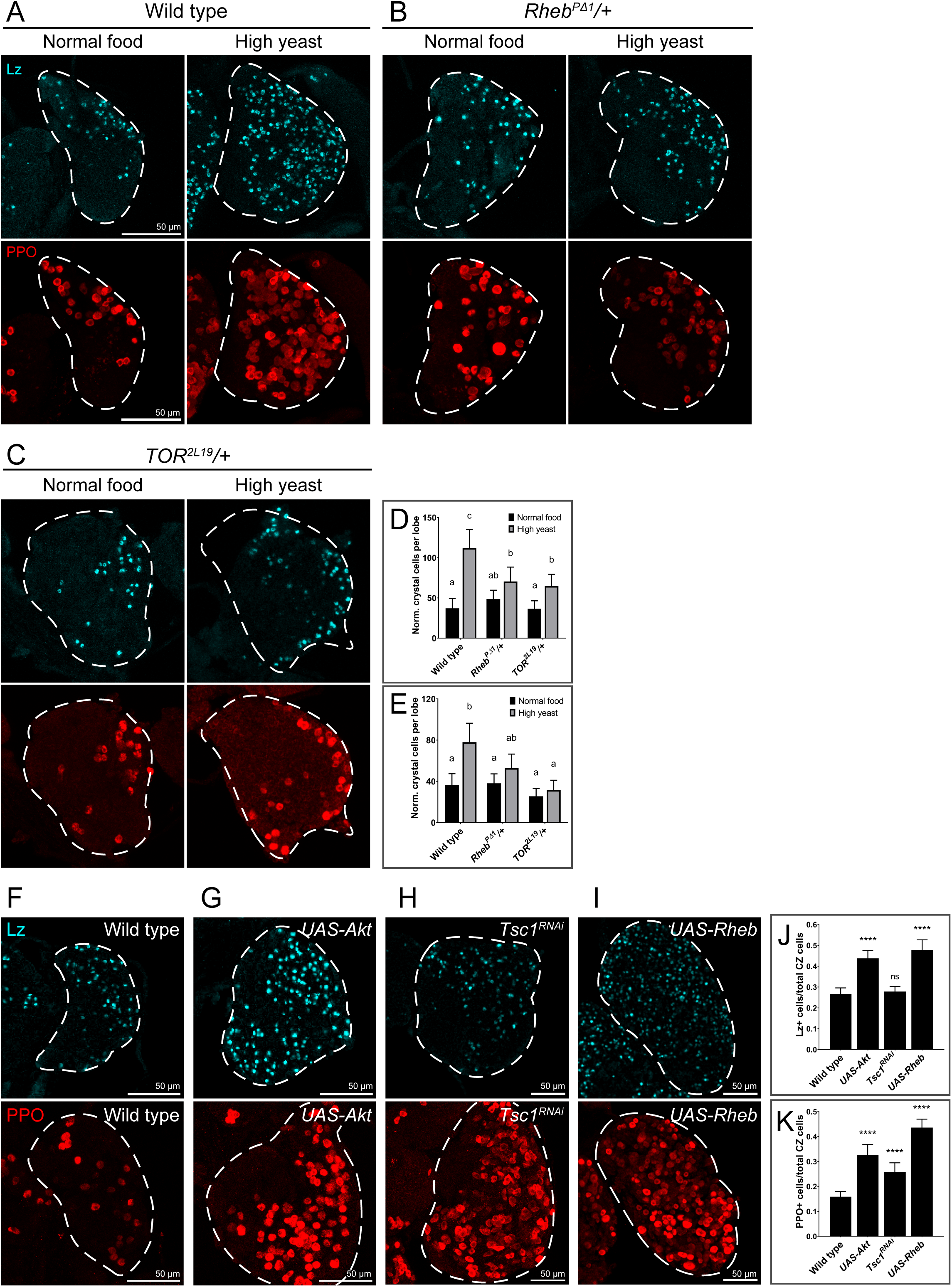
The TOR pathway conveyed the effect of a high yeast diet on crystal cell differentiation. The number of crystal cells was assessed by anti-Lozenge (Lz) and anti-prophenoloxidase (PPO) staining. The increase of crystal cell differentiation induced by a high-yeast diet (A) was mitigated in *Rheb^PΔ1^* (B) or *TOR^2L19^* (C) heterozygous larvae. (D, E) Quantification of the above results, shown as mean crystal cells per lobe ± 95% CI. Two-way ANOVA followed by Tukey’s test for multiple comparisons. Different letters represent significant statistical differences (p<0.05). For the wild type genotype, n = 48 primary lobes in normal food and n = 46 in high yeast; for *Rheb^PΔ1/+^*, n = 13 in normal food and n = 14 in high yeast; for *TOR^2L19/+^*, n = 23 in normal food and n = 23 in high yeast. (F-I) Overexpression of Akt, silencing of Tsc1 or overexpression of Rheb driven by *domeMESO*-Gal4 led to increased crystal cell differentiation. (J, K) Quantification of these results, represented as the ratio between crystal cells and total cortical zone (CZ) hemocytes, which were calculated as the sum of P1+ cells (mature plasmatocytes) and Lz+/PPO+ cells (crystal cells). Mean ± 95% CI. One-way ANOVA followed by Dunnett’s multiple comparisons test (treatments versus control). For wild type, n = 27 for Lz and n = 71 for PPO; for *UAS-Akt*, n = 18 for Lz and n = 16 for PPO; for *Tsc1^RNAi^*, n = 35 for Lz and n = 36 for PPO; for *UAS-Rheb*, n = 22 for Lz and n = 26 for PPO. ****p<0.0001, ns, not significant (p>0.05).

In summary, our results indicate that amino acid abundance can control the extent of crystal cell differentiation in the larval lymph gland by modulating the TOR pathway, which in turn inhibits autophagy. Autophagy inhibition provokes an increase of Notch protein levels, which in turn promotes crystal cell differentiation. Autophagy-dependent regulation of Notch emerges from a crosstalk between the endocytic and autophagy pathways: In conditions of reduced autophagy (high amino acids), Notch lysosomal degradation is hindered, thus favoring Notch accumulation in the limiting membrane of late endosomes/multivesicular bodies, leading to increased Notch signaling and crystal cell differentiation.

## Discussion

In this work, we have shown that autophagy is critically required in blood cell progenitors of the *Drosophila* lymph gland to set a limit to crystal cell differentiation. Autophagy exerts this control by promoting Notch degradation through a crosstalk with the endocytic pathway. As previously reported in other biological settings, we have found that both Notch activation and Notch degradation take place in compartments of the endocytic pathway of blood cell progenitors (Hori et al., 2004; Vaccari et al., 2008; Schneider et al., 2013; Johnson et al., 2016; Shimizu et al., 2024). Upon genetic blockage of endocytosis, or hindering maturation of early endosomes in blood cell progenitors, Notch-dependent differentiation of CCs was impaired. Conversely, by blocking components of ESCRT complexes or of its effector ATPase Vps4, which mediate formation of intraluminal vesicles of the MVB, Notch degradation was impaired, leading to increased CC differentiation. We found that autophagosomes are necessary for Notch degradation. This is consistent with a model in which fusion of MVBs with autophagosomes –giving rise to amphisomes- is necessary for Notch lysosomal degradation. According to this model, autophagy inhibition is expected to reduce Notch lysosomal degradation, leading to increased Notch signaling, and augmented CC differentiation. In agreement with such mechanism, genetic or physiologic overactivation of the TOR pathway, which mediates inhibition of autophagy (Scott et al., 2004; Neufeld, 2010), increased CC differentiation. Thus, our results establish a mechanistic link between the nutritional status of the organism and differentiation of cells of the immune system, in which the Notch pathway is regulated by TOR signaling through the modulation of autophagy.

It is increasingly evident that autophagy is required for proper differentiation or maintenance of most mammalian hematopoietic lineages (Riffelmacher and Simon, 2017; Clarke and Simon, 2019), although the mechanisms involved are incompletely defined. Paralleling our observations in *Drosophila* blood cell progenitors, autophagy is also highly active in mammalian hematopoietic stem cells (Warr et al., 2013), where autophagy is presumably required for lowering reactive oxygen species through mitophagy (Jensen et al., 2011). In T-lymphocytes, as well as in some myeloid lineages, autophagy is believed to control the OXPHOS/glycolytic ratio, thereby impinging on differentiation (Suda et al., 2011; Rožman et al., 2015). During B-cell maturation to antibody-producing plasma cells, autophagy is required to limit the unfolded protein response at the ER, and accumulation of protein aggregates, which in turn has an impact on differentiation (Pengo et al., 2013).

Whereas in *Drosophila* hematopoiesis the primary role of Notch is to promote crystal cell differentiation and maturation (Duvic et al., 2002; Mukherjee et al., 2011; Terriente-Felix et al., 2013; Ferguson and Martinez-Agosto, 2014; Blanco-Obregon et al., 2020; Ho et al., 2023), in mammalian hematopoiesis Notch has a more complex, and incompletely understood function. Notch has been reported to act either as a tumor suppressor or as a tumor promoter in different types of leukemia (Kushwah et al., 2014; Láinez-González et al., 2022), and consistent with this, Notch fulfills different roles depending on the hematopoietic lineage (Stier et al., 2002; Burns et al., 2005; Lee et al., 2008b; Mercher et al., 2008; Wang et al., 2014b; Suresh and Irvine, 2015).

TOR dependent regulation of Notch was previously reported in other *Drosophila* organs, even though the mechanisms involved were not completely defined. In *Drosophila* enteroblasts, regulation of Notch by the TOR pathway was shown to depend on miRNAs (Foronda et al., 2014), which does not exclude the possibility that TOR might regulate Notch through an autophagy-dependent mechanism in the *Drosophila* gut as well. Autophagy-dependent degradation of Notch apparently takes place in *Drosophila* follicle cells by mid-oogenesis (Barth et al., 2012) opening the possibility that the complete mechanism described here may be more general and apply to other *Drosophila* organs in addition to hematopoietic progenitors.

Evidence for autophagy-dependent destruction of Notch has also been reported in mammalian cells (Wu et al., 2016), although the mechanism in that context involves direct uptake of Notch into Atg16L1-positive vesicles, rather than into regular Rab5/Rab7-positive endosomes. It remains to be assessed if the cellular mechanisms described here in operate in mammalian systems as well.

Summing up, the mechanism described here for autophagy-mediated regulation of Notch pathway activity in *Drosophila* blood cell progenitors might be active in other organs and model systems as well. Further research is therefore required to determine to what extent other Notch-dependent differentiation events are controlled by nutrient availability through the TOR-autophagy axis.

## Materials and Methods

Flies were reared at 25°C on a standard cornmeal/agar medium. For Gal4-mediated genetic manipulations, all experimental crosses were conducted at 25°C, with F1 first instar larvae subsequently incubated at 29°C to maximize Gal4 activity. To assess the nutritional impact on CC differentiation, standard cornmeal medium supplemented with 4% or 2% yeast was utilized. The following fly stocks were obtained from the Bloomington *Drosophila* Stock Center (https://bdsc.indiana.edu): *w^1118^* (BL 3605), Canton-S (BL 9515), *UAS-GFP* (BL 1521), *YFP-Rab7* (BL 62545), *YFP-Rab5* (BL 52543), *Notch^55e11^* (BL 28813), *Su(H)^1^* (BL 417), *Atg1^RNAi^* (BL 26731), *Atg17^RNAi^* (BL 36918), *Atg18^RNAi^* (BL 34714), *Vps15^RNAi^* (BL 34092), *Vps34^RNAi^* (BL 33384), *Notch^RNAi^* (BL 33611), *Su(dx)^RNAi^* (BL 67012), *E(spl)mß-HLH-GFP* (BL 65294), *shibire^RNAi^* (BL 28513), *Hrs^RNAi^* (BL 33900), *UAS-Rab5^DN^* (BL 9772), *Tsg101^RNAi^* (BL 38306), *Vps25^RNAi^* (BL 54831), *Fab1^RNAi^* (BL 35793), *Vamp7^RNAi^* (BL 38300), *Syx17^RNAi^* (BL 29546), *UAS-Atg1* (BL 60734), *slimfast^RNAi^* (BL 64922), *UAS-slimfast* (BL 52661), *UAS-Akt* (BL 8191), *Tsc1^RNAi^* (BL 52931) and *UAS-Rheb* (BL 9688). *shrub^RNAi^* (v108557) and *Vps4^RNAi^* (v105917) were obtained from the Vienna *Drosophila* Resource Center (https://stockcenter.vdrc.at). The following stocks were kindly provided by colleagues: *domeMESO-*Gal4 (Utpal Banerjee), *3xmCherry-Atg8a* and *Atg1*^Δ3D^ (Gábor Juhász), *UAS-LacZ* (Ben Shilo), *UAS-GFP-Ref2P* (Thomas Neufeld), *UAS-GFP-Lamp* (Helmut Kramer), *UAS-deltex* (Spyros Artavanis-Tsakonas), *Rheb^PΔ1^* and *TOR^2L19^* (Sean Oldham).

### Dissection, fixation and immunostaining of lymph glands

Lymph glands were processed and stained as previously described (Evans et al., 2014). Briefly, wandering third instar larvae were used for dissection of lymph glands in PBS. Dissected tissue was fixed in 4% formaldehyde for 30 minutes and washed three times in PBS with 0.4% Triton-X (0.4% PBST) for 15 minutes. Blocking was carried out in 10% normal goat serum (Sigma-Aldrich) in PBST for 30 minutes, followed by overnight incubation with primary antibodies at 4°C in 10% normal goat serum. Thereafter, lymph glands were washed three times in 0.4% PBST for 15 minutes, and incubated with secondary antibodies for 1.5 h at room temperature in 10% normal goat serum. Finally, samples were washed three times in 0.4% PBST before mounting on glass slides in mounting medium (Mowiol 4-88 anti-fade agent, EMD Millipore Corp., Billerica, MA). The following primary antibodies were used: mouse α-Lozenge 1:10 (anti-lozenge, Developmental Studies Hybridoma Bank), mouse α-Notch Extracellular Domain 1:10 (C458.2H, Developmental Studies Hybridoma Bank), mouse α-P1 1:100 (gift from I. Ando), rabbit α-PPO 1:2000 (generous gift from G. Christophides). Alexa Fluor 488, Alexa Fluor 647 and Cy3-conjugated secondary antibodies 115-545-062, 115-605-166, and 111-165-144, respectively, Jackson ImmunoResearch) were utilized in 1:250 dilutions.

### Image Acquisition and Processing

Lymph glands were imaged using a Zeiss LSM 880 confocal microscope, acquiring either Z-stacks or single confocal planes. Image processing was conducted using Fiji/ImageJ software (NIH). Lobe area quantification was performed by manually delimiting lobe perimeter. *domeMESO* and P1 areas were automatically calculated. For crystal cell quantification, the total number of CCs was quantified automatically based on nuclear markers such as Lz and E(spl)-GFP, and manually when using the cytoplasmic marker PPO, then corrected for lobe area in comparison with the control treatment whenever a significant difference in lobe size was found between the treatments (whole Z-stack projections). In experiments where a notorious change in cell size, as well as in lobe size, was observed (Fig 9F-K), crystal cell differentiation was assessed by calculating the ratio of crystal cells to differentiated hemocytes (P1-positive for plasmatocytes and Lz-positive or PPO-positive for crystal cells) on a representative slice of the Z-stack. Notch levels were calculated as mean intensity of the whole primary lobe, except for Notch intensity measure in MVBs (SFig 5F), where quantification was restricted to Rab7-positive area. 3xmCherry-Atg8a, GFP-Ref(2)P and Lysotracker *foci* were automatically quantified.

### Statistical analysis

Statistical analysis was performed on GraphPad Prism version 8 and RStudio (R version 4.2.3). The threshold for statistical significance was established as p<0.05. Each experiment was independently repeated at least twice.

## Acknowledgements

We are grateful to many colleagues of the *Drosophila* community, the Bloomington Stock Center and the Vienna *Drosophila* Resource Centre for fly strains. Andrés Rossi and Esteban Miglietta, from FIL microscopy facility, for technical support; Andrés Liceri for fly food preparation; the FIL personnel for assistance and members of the Wappner lab for fruitful discussions. This work was supported by grants from Agencia Nacional de Promoción de Científica y Tecnológica: PICT-2018-1501 and PICT-2021-I-A-00240 to PW and PICT 2019-0621 and PICT 2015-0225 to MJK. FR was supported with a fellowship of the ANPCyT and FE with a fellowship of the Consejo Nacional de Investigaciones Científicas y Técnicas.

## Supplementary Figure Legends

**Supplementary Figure 1- Autophagy activation in various *Drosophila* larval organs.** (A-E) The extent of autophagy activation was analyzed in the indicated larval organs by the nucleation of the 3xmCherry-Atg8a autophagy reporter, and by the nucleation of the Lysotracker-green reporter. Quantification of the results in (F) and (G) respectively, showing mean ± 95% CI. For 3xmCherry-Atg8a, n ≥ 8, for Lysotracker, n ≥ 5. One-way ANOVA followed by Dunnett’s multiple comparisons test. Autophagy is induced by starvation in the larval lymph gland, as assessed by the nucleation of the 3xmCherry-Atg8a reporter (H), the Lysotracker-green reporter (I), and by fluorescence intensity of the GFP-Ref(2)P autophagy flux reporter (J). Quantification of the above results is shown as mean ± 95% CI. For 3xmCherry-Atg8a, n ≥ 4 primary lobes; for Lysotracker, n ≥ 8; for GFP-Ref(2)P, n ≥ 4. Student’s *t*-tests. **p<0.01, ***p<0.001, ****p<0.0001.

**Supplementary Figure 2- Autophagy activation in the larval lymph gland depends on canonical autophagy genes.** The 3xmCherry-Atg8a autophagy reporter and the autophagy flux reporter GFP-Ref(2)P were utilized to assess autophagy activation in larvae expressing the indicated interference RNAs (A-F) under control of a *domeMESO*-Gal4 driver that mediates expression in lymph gland progenitor cells. Autophagy is significantly reduced after silencing of any of the canonical autophagy genes. (G, H) Quantification of the results of 3xmCherry-ATG8a nucleation, and GFP-Ref(2)P fluorescence intensity respectively. Mean ± 95% CI. (G) For wild type, n = 29 primary lobes; for *Atg1^RNAi^,* n = 19; for *Atg17^RNAi^*, n = 22; for *Vps15^RNAi^*, n = 20; for *Vps34^RNAi^*, n = 17; for *Atg18^RNAi^*, n = 10. One-way ANOVA followed by Dunnett’s multiple comparisons test. (H) For wild type, n = 33; for *Atg1^RNAi^,* n = 9; for *Atg17^RNAi^*, n = 7; for *Vps15^RNAi^*, n = 22; for *Vps34^RNAi^*, n = 12; for *Atg18^RNAi^*, n = 31. Kruskal-Wallis test followed by Dunn’s multiple comparisons test. ****p<0.0001, ns, not significant (p>0.05).

**Supplementary Figure 3- Activation of the Notch ligand-independent pathway induced crystal cell differentiation.** (A-C) Differentiation of crystal cells in the lymph gland was assessed by anti-Lozenge immunofluorescence (Lz). Overexpression of deltex (B), or silencing of Suppressor of deltex (C) provoked increased crystal cell differentiation. (D) Quantification of the results of panels (A-C), depicting the mean ± 95% CI. For wild type, n = 22 primary lobes; for *UAS-deltex,* n = 52; for *Su(dx)^RNAi^*, n = 17. Likelihood Ratio Test followed by Dunnett’s multiple comparisons test. ****p<0.0001.

**Supplementary Figure 4- Antagonistic genetic interactions between the autophagy pathway and genes of the Notch ligand-independent pathway.** (A-F) Crystal cell differentiation was evaluated by immunofluorescence with an anti-prophenoloxidase antibody (PPO). Knock-down of deltex (dx) (C) or overexpression of Suppressor of deltex (Su(dx)) (E) did not modify the number of crystal cells. Knock-down of Atg1 (B) provoked increased crystal cell differentiation, but this increase was eliminated when deltex was knocked-down (D), or Suppressor of deltex was overexpressed (F) with a *domeMESO*-Gal4 driver. (G) Quantification of the results of panels (A-F), representing mean ± 95% CI. For wild type, n = 22 primary lobes; for *Atg1^RNAi^,* n = 20; for *dx^RNAi^*, n = 23; for *dx^RNA^/Atg1^RNAi^*, n = 21; for *UAS-Su(dx)*, n = 23; for *UAS-Su(dx); Atg1^RNAi^*, n = 20. One-way ANOVA followed by Tukey’s multiple comparisons test. Different letters account for significant statistical differences (p<0.05).

**Supplementary Figure 5- Notch localized at endosomal compartments where accumulated after autophagy inhibition.** (A) Schematic representation of endosomal maturation and fusion with autophagosomes. Endosomes with the Notch Intracellular Domain (NICD) inserted in their membranes mature into multivesicular bodies, in which limiting membrane the gamma-secretase complex (γ-sec) can cleave and release the NICD that enters the nucleus (nucleus not shown in the scheme). Alternatively, invagination of the limiting membrane of the multivesicular body occurs, giving rise to intraluminal vesicles of the multivesicular body that eventually contain the membrane-bound NICD. Multivesicular bodies then fuse with autophagosomes to give rise to NICD-containing amphisomes, which finally fuse with lysosomes, resulting in autolysosomes where destruction of the NICD takes place. (B-D) High-resolution (AiryScan) confocal images of Notch associated to endosomal vesicles. (B) Part of Notch protein (cyan, white arrowheads) localized in Rab5-positive early endosomes (red, yellow arrowheads). (C) Part of Notch protein (cyan, white arrowheads) localized in Rab7-positive late endosomes/multivesicular bodies (red, yellow arrowheads). (D) After autophagy inhibition through the expression of *Atg1^RNAi^*, Rab7-positive vesicles significantly enlarged (red, yellow arrowheads) and contained increased amounts of Notch protein (cyan, white arrowheads). (E) Quantification of the size of Rab7-positive vesicles in the experiments of panels (C, D) and after silencing the indicated autophagy genes, expressed as mean ± 95% CI. n ≥ 628 vesicles for each genotype. (F) Quantification of Notch in Rab7 vesicles in the experiments of panels (C, D), and after silencing of the indicated autophagy pathway genes. Data are expressed as relative change of cyan fluorescence intensity associated with Rab7 vesicles (red). Mean ± 95% CI, n ≥ 966 vesicles for each genotype. The statistical analysis in both cases was Kruskal-Wallis test followed by Dunn’s multiple comparisons test (treatments vs. control). **p<0.01, ****p<0.0001.

**Supplementary Figure 6- Notch colocalized with autophagosomal, amphisomal and lysosomal markers.** Super-resolution (Airyscan) confocal images of Notch associated to autophagosomal and lysosomal structures. (A) Notch (cyan, white arrowheads) was observed colocalizing with Rab-7 positive vesicles (green, yellow arrowhead), and with Atg8a-positive structures (red, pink arrowheads). (B) Notch (cyan, white arrowheads) appeared associated with Lamp positive vesicles (green, yellow arrowheads) that also contained Atg8a (red, pink arrowheads).

**Supplementary Figure 7- Autophagy was reduced in yeast-rich larval culture medium or upon genetic overactivation of the TOR pathway.** Autophagy activation was assessed by the nucleation of the 3xmCherry-Atg8a reporter. (A) Nucleation of the reporter is higher in culture medium containing 2% yeast (w/v) (normal food), as compared to medium with 4% yeast (high yeast). (B) Quantification of the results depicted in (A). Mean ± 95% CI, n = 22 primary lobes for normal food medium, n = 30 for high yeast medium. Student’s *t*-test. (D-G) Autophagy activation was strongly reduced after expression of the indicated transgenes that activate TOR signalling. A *domeMESO*-Gal4 driver was utilized to induce expression in blood cell progenitors. (H) Quantification of the autophagy activation in the experiments of panels (D-G). Mean ± 95% CI. n = 47 primary lobes for wild type; n = 18 for *UAS-Akt*; n = 16 for *Tsc1^RNAi^*; n = 25 for *UAS-Rheb*. Kruskal-Wallis test followed by Dunn’s multiple comparisons test. ***p<0.001, ****p<0.0001.

