## Supplementary figures and images for "Autophagy controls differentiation of *Drosophila* blood cells by regulating Notch levels in response to nutrient availability"

### Supplementary Figure 1

#### Supplementary Figure 1

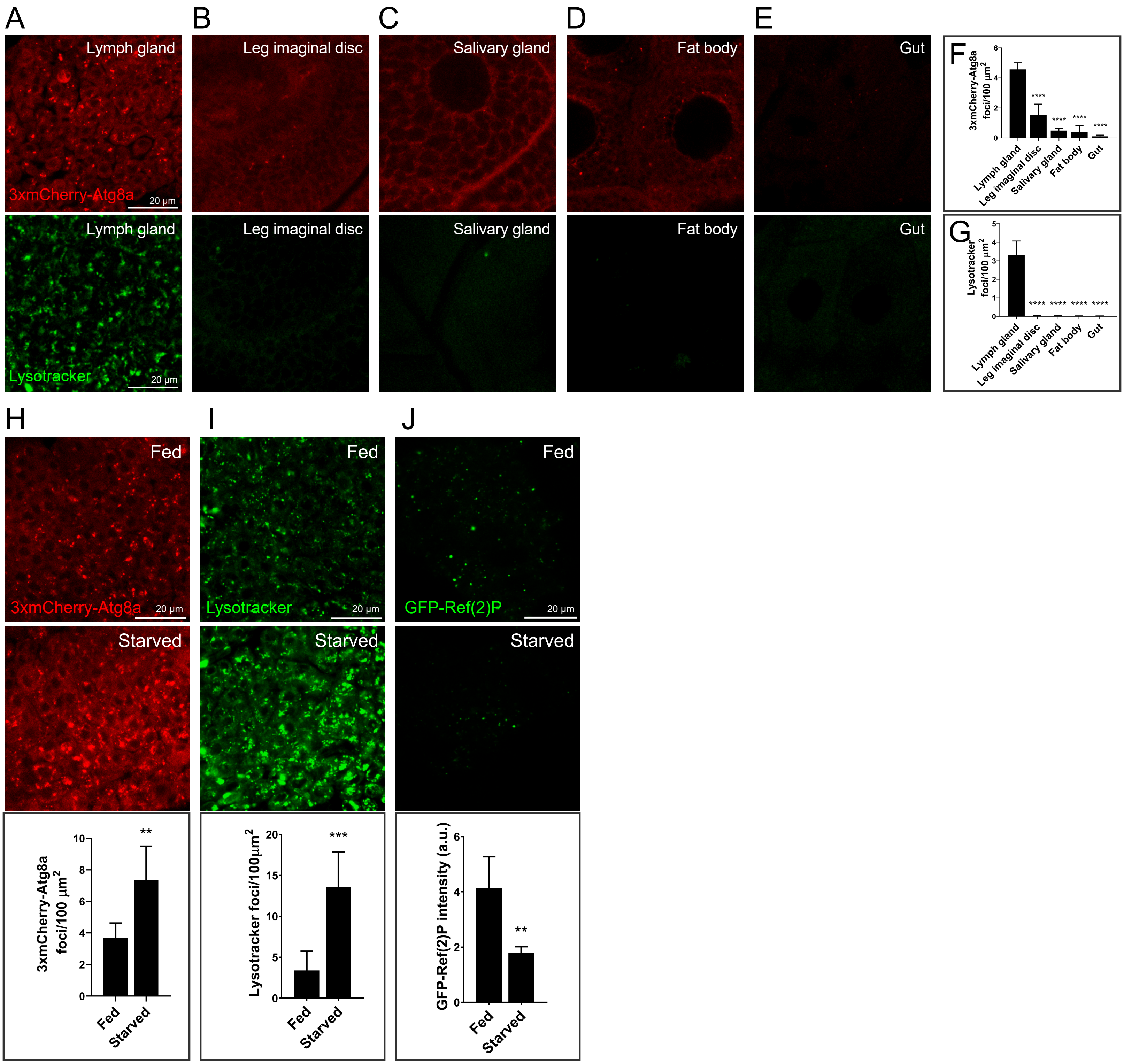

### Supplementary Figure 2

#### Supplementary Figure 2

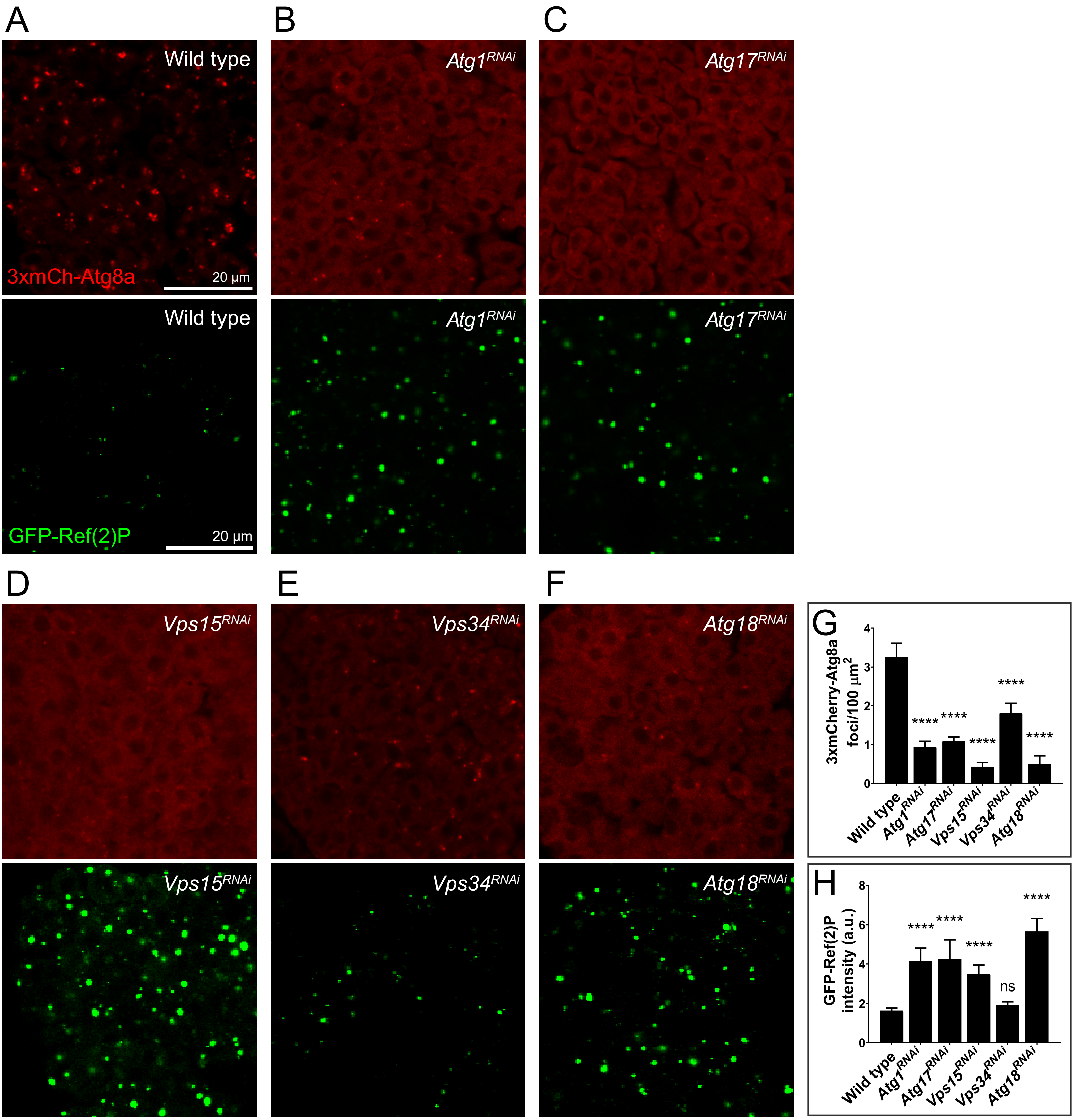

### Supplementary Figure 3

Supplementary Figure 3

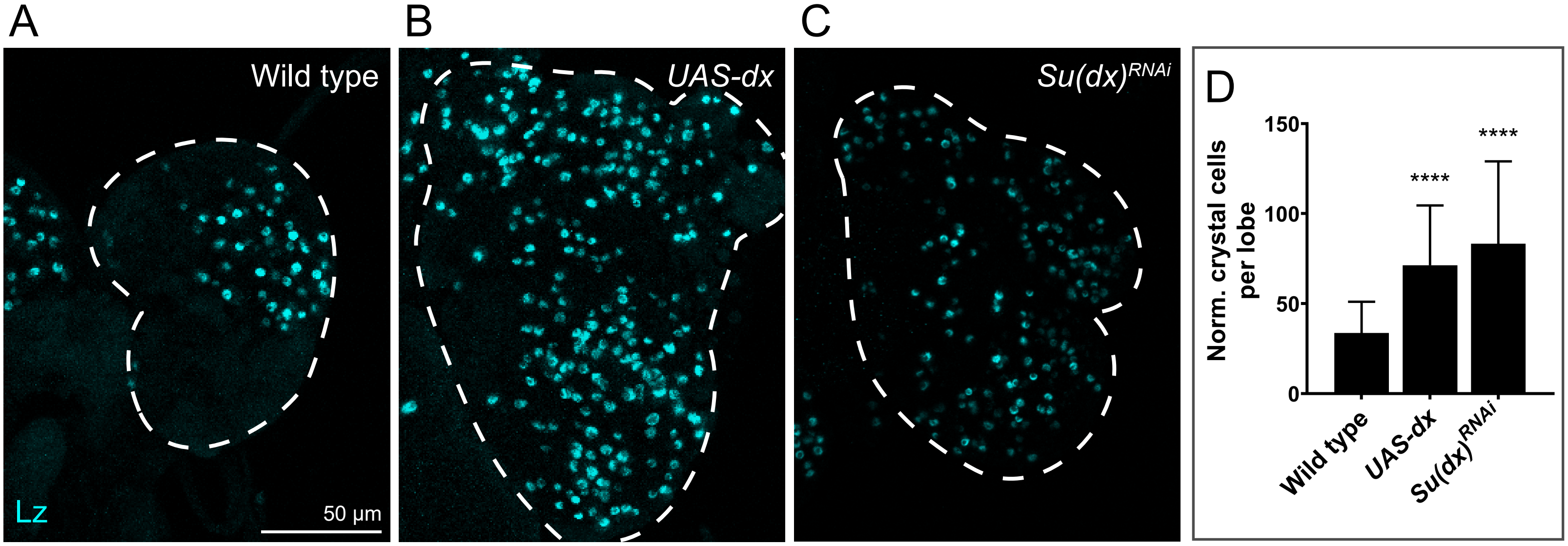

### Supplementary Figure 4

Supplementary Figure 4

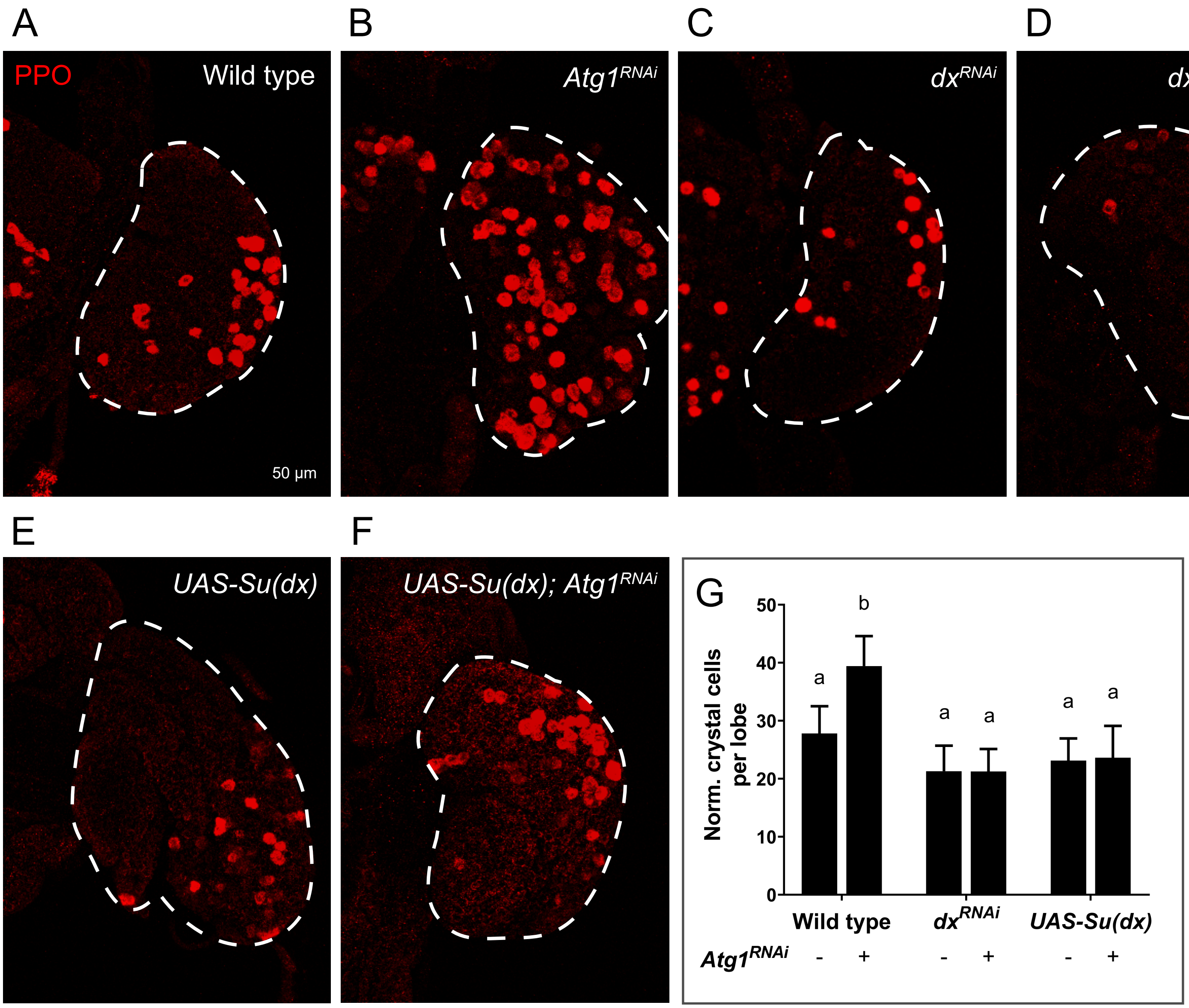

### Supplementary Figure 5

Supplementary Figure 5

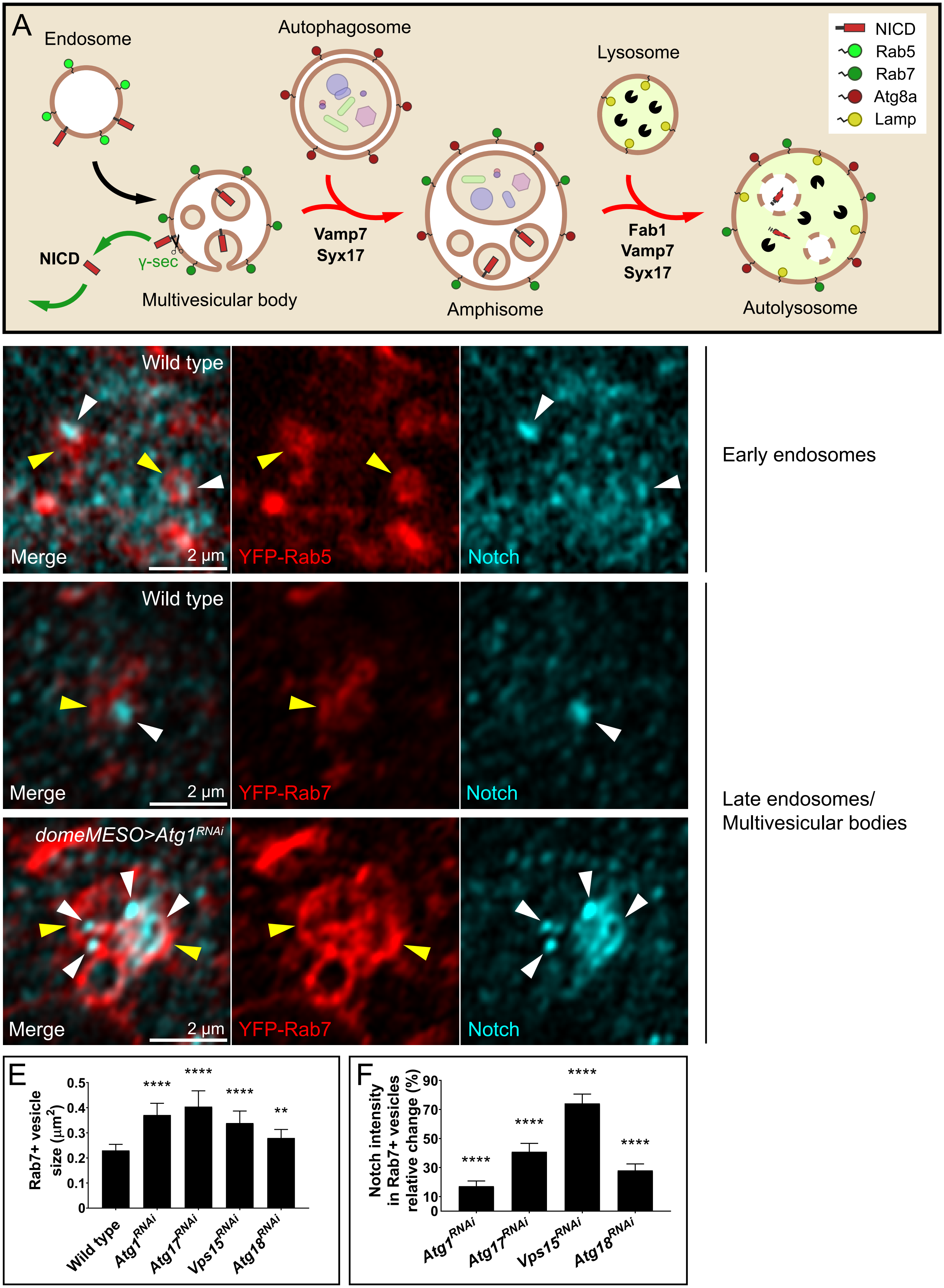

### Supplementary Figure 6

Supplementary Figure 6

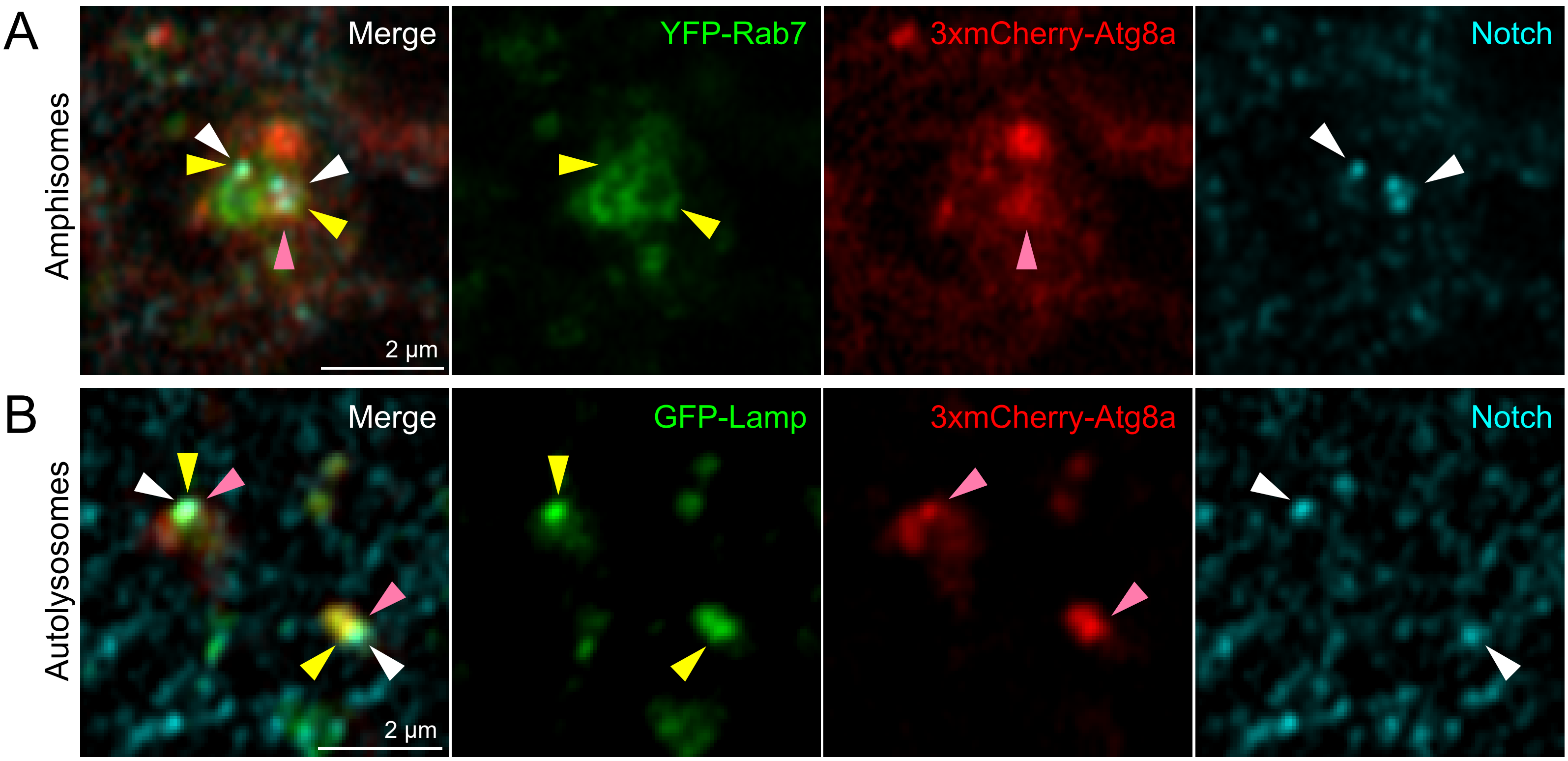

### Supplementary Figure 7

#### Supplementary Figure 7

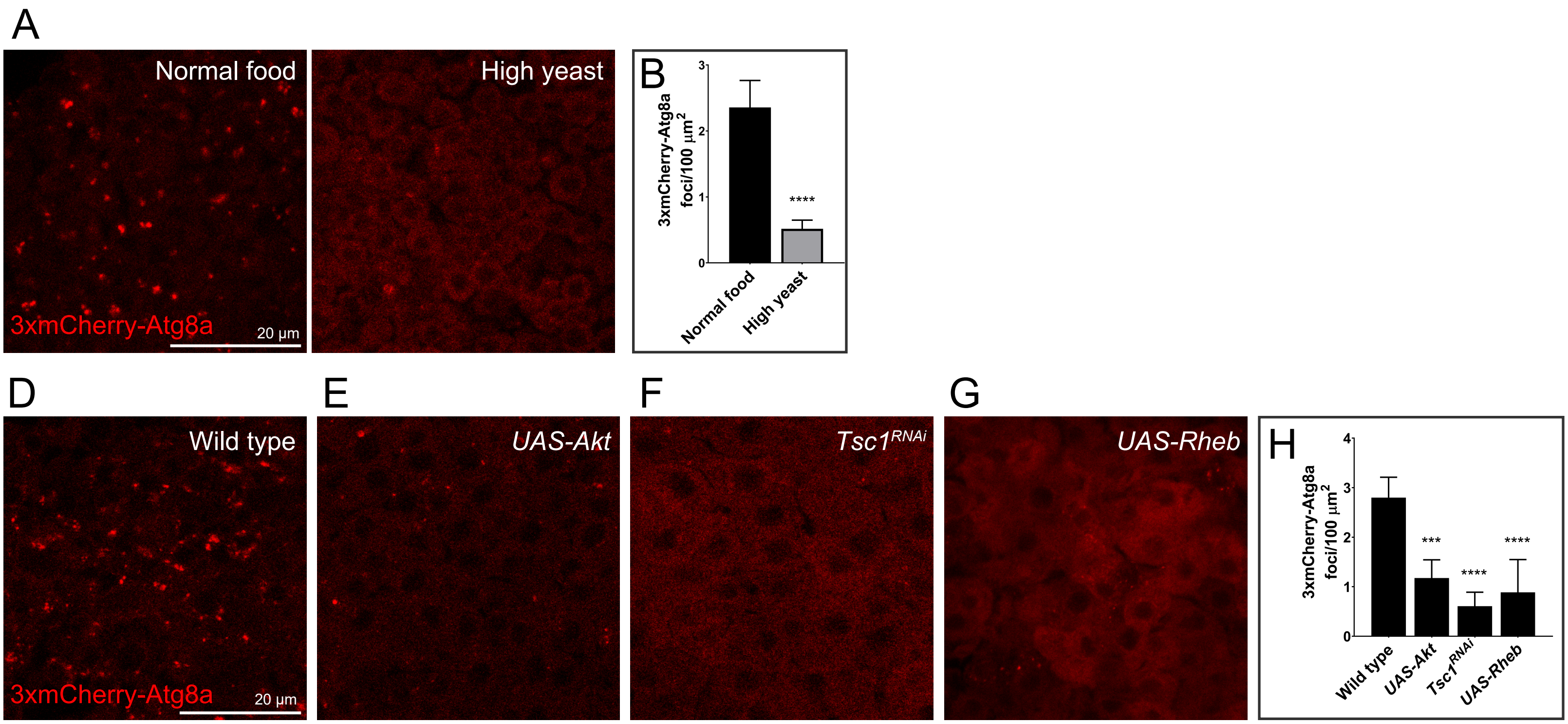
